# The translocation activity of Rad54 reduces crossover outcomes during homologous recombination

**DOI:** 10.1101/2024.01.25.577253

**Authors:** Krishay Sridalla, Mitchell V. Woodhouse, Jingyi Hu, Jessica Scheer, Bryan Ferlez, J. Brooks Crickard

## Abstract

Homologous recombination (HR) is a template-based DNA double-strand break repair pathway that requires the selection of an appropriate DNA template for repair during the homology search stage of HR. Failure to execute the homology search quickly and efficiently can result in complex intermediates that generate genomic rearrangements, a hallmark of human cancers. Rad54 is an ATP dependent DNA motor protein that functions during the homology search by regulating the recombinase Rad51. How this regulation reduces genomic rearrangements is currently unknown. To better understand how Rad54 can prevent genomic rearrangements, we evaluated several amino acid mutations in Rad54 that were found in the COSMIC database. COSMIC is a collection of amino acid mutations identified in human cancers. These substitutions led to reduced Rad54 function and the discovery of a conserved motif in Rad54. Through genetic, biochemical, and single-molecule approaches, we show that disruption of this motif leads to failure in stabilizing early strand invasion intermediates, causing loss-of-heterozygosity rearrangements. Our study also suggests that the translocation rate of Rad54 is a determinant in balancing genetic exchange. This mechanism is likely fundamental to eukaryotic biology.

## Introduction

Homologous recombination (HR) is a template-based DNA double strand break repair (DSBR) pathway based on locating a matching sequence element elsewhere in the genome (1–4). To find a DNA template, HR proceeds through a series of steps that begins with the identification of a double-strand break followed by a 5’ to 3’ resection of the DNA to create ssDNA overhangs (5, 6). The ssDNA is then loaded with recombinase filaments, Rad51 in eukaryotes, which use the ssDNA as a guide to search the genome for a template to facilitate DNA repair (7–11). Once a template is found, Rad51, in conjunction with numerous accessory factors, promotes a strand invasion reaction, which results in further base pairing between the recipient ssDNA and the donor dsDNA as well as displacement of the non-template strand, resulting in the formation of a structure known as the displacement loop (D-loop) (12–15).

Formation of the D-loop during HR is a dynamic process that requires the selection of an appropriate template and stabilization of strand pairing events in a process called the homology search (7–11, 16–22). The end point of the homology search is the extension of D-loops, which allows the transition to the DNA repair and resolution phase of HR (23). In *Saccharomyces cerevisiae,* the steps in the homology search are regulated by several DNA translocases and helicases, including Rad54 (24), Rdh54 (14, 25–27), and Srs2 (13, 28–32). These ATP-dependent motors control intermediates, including Rad51 filaments (28, 29, 32, 33) and D-loop structures (13, 33, 34). Each of these motors also impacts HR outcomes with the deletion of RDH54, resulting in increased break-induced replication (BIR), and deletion of SRS2, resulting in increased DNA crossovers (CO) between ectopic DNA sequences (35). Importantly, increases in these outcomes can have mutagenic consequences on the genome. The role of Rad54 in preventing genomic re-arrangements is poorly defined, given its general importance to D-loop formation. Thus, making it difficult to disentangle early steps from late steps.

Rad54 is a Snf2 family DNA motor protein that functions as a multimeric motor during the homology search to promote DNA and chromatin remodeling (36–39), Rad51 remodeling (40–42), and D-loop stabilization (15, 34, 40, 43). Deletion of Rad54 creates a reduction in D-loop formation, leading to failure in HR mediated repair (34) causing chromosome loss (44). Genetic evidence suggests that overexpression of catalytically inactive Rad54 presents an increased crossover between homologous chromosomes phenotype, a type of genomic rearrangement in mitotic cells. A dominant negative effect could cause this as Rad54 functions as a multimer. This also infers a disruption in the synthesis-dependent strand annealing pathway (SDSA), generally accepted to promote non-crossover outcomes during DNA repair (1, 2, 7, 45). This observation is consistent with the activity of Rad54 being required to prevent complex chromosomal rearrangement events (46). The mechanism by which Rad54 prevents excessive strand exchange is currently unclear. However, Rad54 is known to promote branch migration, and it is speculated that this activity promotes SDSA and reduces the extent of DNA exchange between donor and recipient DNA (47–49).

The roles of Rad54 during HR have led to the development of the idea that this motor acts as a general catalyst to speed up the DNA repair reaction. This is consistent with the model that Rad54 works as a sister chromatid exchange factor (50, 51), as accelerating HR will likely favor interactions between sister chromatids. Failure to initiate rapid repair can lead to increased DNA end resection (52, 53) and the accumulation of ssDNA within the genome, leading to complex rearrangements (54, 55). Complex chromosomal rearrangements within the genome characterize many human cancers due to defects in DSBR pathways. Re-arrangements can also lead to the development and the evolution of existing tumors (56–59). To better define the role of Rad54 in mitigating genomic rearrangements, we sought to identify and characterize amino acid substitutions in Rad54 from human cancers that slowed down its translocation activity. Our data suggests that reduction in the Rad54 translocation activity leads to a kinetic delay in strand invasion, which can result in genomic re-arrangements characterized as crossovers between homologous chromosomes. We develop a model from which factors that stabilize early strand invasion intermediates may kinetically prevent complex rearrangements within the genome.

## Materials and Methods

### Yeast strain construction

All recombination outcome experiments were performed in the W303 background. Strains for spot assay experiments were BY4741 and were generated by transforming *rad54*Δ strains with pRS415 plasmids. For integrated spot assays, *rad54* mutations were introduced by gene replacement into BY4741 strains. The genotypes for all strains used in this study can be found in **Supplemental Table 1**. Plasmids for the generation of strains can be found in **Supplemental Table 2**.

### Yeast spot growth assay and colony growth

For complementation spot assays, overnight cultures were diluted back to an OD_600_ of 0.2 and then allowed to grow to an OD_600_ of 1.0. Cells were then serially diluted and manually spotted on YNB -Leu +2% Dextrose plates containing no drug, 0.003 or 0.01% methylmethanesulfonate (MMS). Plates were incubated at 30°C for 2-3 days and imaged at 24, 48, and 72 hours. The same protocol was followed for integrated constructs, except the cells were grown and spotted on YPD plates. In the case of Rad51 overexpression experiments. pYes-RAD51 plasmids were transformed into strains with *rad54R272Q* and *rad54R272A* alleles integrated into the genome. Spot assays were performed as described above, except cultures were spotted on YNB (-Ura) +/−2% dextrose or YNB+/− (-Ura) 2% galactose. The plates were grown for 48 and 72 hours and imaged.

### Red/White recombination assay

The WT strain used in this assay and the procedure for diploid formation are described here (27). The genotypes for modifying these strains can be found in **Supplemental Table 2**. The assay was performed by growing the appropriate strain overnight in YP + 2% raffinose. The next day, cells were diluted to an OD600 of 0.2 and allowed to reach an OD600 of 0.4-0.5, followed by the expression of *I-Sce1* by adding 2% galactose. Cells were allowed to grow for an additional 1.5 hours, then plated on YPD plates, and allowed to grow for 48 hours. After 48 hours, they were placed at 4 °C overnight to enable further development of the red color. The number of white, red, and sectored colonies was then counted followed by replica plating onto YPD + hygromycin B (200 µg/ml) and YPD + nourseothricin (67 µg/ml, clonNat) for analysis of recombination outcomes. Strains were also replica plated on YNB (-Ura/-Met) + 2% dextrose to ensure proper chromosome segregation (**Supplemental Table 4-6**). The data was analyzed by counting sectored colonies and colony survival on different antibiotic sensitivities. The data for each category was then divided by the total population of sectored colonies. The standard deviation between biological replicates was analyzed for at least three independent experiments from different crosses.

### Protein Purification

Rad51, Rad54, and RPA purifications were performed as previously described (8).

### Electrophoresis mobility shift assay (EMSA) for Rad54

An Atto647N labeled 90-mer oligo was annealed with an unlabeled complementary oligo to make a labeled dsDNA substrate. The sequences of the oligos used in this assay can be found in **Supplemental Table 7.** The assay was performed in EMSA buffer (35 mM Tris-Cl pH 7.5, 3 mM MgCl_2_, 50 mM KCl, 1 mM DTT, 10% Glycerol). The final DNA concentration was 10 nM and proteins were titrated to be 0, 6.25, 12.5, 25 and 50 nM as final concentrations. The DNA and proteins were incubated at 30 °C for 5 minutes and then resolved by 10% Native-PAGE (20 mM Tris, 10 mM acetic acid, 0.5 mM EDTA, 10% acrylamide/bis-acrylamide (37.5:1), 0.1% APS, 0.1% TEMED) and ran in 0.5x TAE buffer (20 mM Tris, 10 mM acetic acid, 0.5 mM EDTA).

### ATPase assay

A commercially available ADP-GLO kit was used to measure ATP hydrolysis rates. The ATP hydrolysis reaction was performed in HR buffer (20 mM Tris-OAc, 50 mM NaCl, 10 mM MgOAc_2_, 200 ng/µl BSA, 1 mM DTT, and 10% Glycerol) and contained 1 mg/ml sheared salmon sperm DNA and 10 nM Rad54.

### *In vitro* D-loop assay

D-loop formation experiments were performed in HR buffer (30 mM Tris-OAc [pH 7.5], 50 mM NaCl, 10 mM MgOAc_2_, 1 mM DTT, 0.2 mg/ml BSA) using an Atto647N labeled DNA duplex (15 nM) with homology to the pUC19 plasmid. Rad51 (300 nM) was incubated with recipient DNA at 30 °C for 15 minutes. The resulting Rad51 filaments were mixed with indicated concentrations of Rad54, RPA (500 nM), and pUC19 plasmid (0.3 nM). Reactions were quenched after indicated periods and treated with 1 unit of Proteinase K at 37°C for 20 minutes. The reactions were then resolved by electrophoresis on a 0.9% agarose gel and imaged for fluorescence using a Typhoon imager. The sequences of the oligos used in this study can be found in **Supplemental Table 7.**

### Flow cell construction

Metallic chrome patterns were deposited on quartz microscope slides with pre-drilled holes for microfluidic line attachment by electron beam lithography to generate flow cells. After metal deposition, a channel was created by covering the two-sided tape with a small piece of paper between the drill holes. The paper was excised to make the flow chamber, and a glass coverslip was fixed to the tape. The chamber was sealed by heating to 165°C in a vacuum oven at 25 mm Hg for 60 minutes. Flow cells were then completed using hot glue to fit IDEX nano ports over the drill holes on the opposite side of the microscope slide from the coverslip.

### Single molecule experiments

All single molecule experiments were conducted on a custom-built prism-based total internal reflection microscope (Nikon) equipped with a 488-nm laser (Coherent Sapphire, 100 mW), a 561-nm laser (Coherent Sapphire, 100 mW), a 640-nm laser (Coherent Obis, 100 mW) and two Andor iXon EMCCD cameras. DNA substrates for DNA curtains experiments were made by attaching a biotinylated oligo to one end of the 50 kb Lambda phage genome and an oligo with a digoxygenin moiety on the other. This allowed double tethering of the DNA between chrome barriers and chrome pedestals, as previously described (60, 61). Flow cells were attached to a microfluidic system, and sample delivery was controlled using a syringe pump (KD Scientific). Three-color imaging was achieved by two XION 512×512 back-thinned Andor EM-CCD cameras and alternative illumination using a 488-nm laser, a 561 nM laser, and a 640 nM laser at 25% power output. The lasers were shuttered resulting in a 200 msec delay between each frame. Images were collected with a 200 msec integration time. Translocation velocity and distances were measured in HR Buffer (20 mM Tris-OAc, 50 mM NaCl, 10 mM MgOAc_2_, 200 ng/µl BSA, 1 mM DTT).

### Analysis of dsDNA translocation

The velocity and track length for GFP-Rad54 molecules were measured by importing raw TIFF images as image stacks into ImageJ. Kymographs were generated by defining a 1-pixel wide region of interest (ROI) along the long axis of individual dsDNA molecules. Data analysis was performed from the kymographs. The start of translocation was defined when the GFP-Rad54 molecule moved > 2 pixels. Pauses were defined as momentary stalls in translocation that lasted 2-4 frames. Termination was defined by molecules that did not move for >10 frames. Velocities were calculated using the following formula [(|Y_f_-Y_i_|)*1,000 bp/[|X_f_-X_i_|])*frame rate]; where Y_i_ and Y_f_ correspond to the initial and final pixel position and X_i_ and X_f_ correspond to the start and stop time (in seconds). Graphs of individual velocity and distances traveled were plotted in GraphPad Prism 9. The mean was determined from these graphs, and significant differences between mutants were determined by performing a student’s t-test of the data. The selection of homologous sequence was performed by measuring the position of GFP-Rad54 and 90-mer Atto647N after 15 minutes. The probability of all bound sequences was then calculated and plotted. A non-homologous piece of DNA was subjected to the same experiments and analyzed as a negative control. Determination of homology search type was performed by visual inspection of kymographs for evidence of translocation before stabilization at the homologous site.

### Single Molecule Analysis of RPA

Analysis of RPA was performed by visually inspecting kymographs for RPA-mCherry signal that colocalized with GFP-Rad54 and Atto647N-ssDNA. The association time for RPA was determined by measuring the number of frames it took for RPA to colocalize with GFP-Rad54 and Atto647N-ssDNA. This number was converted to seconds, and then populations were compared. For lifetime measurements, the number of frames with a detectable RPA signal was scored and converted to seconds. This was then plotted on an exponential decay curve to determine the percentage dissociation. The dissociation rate was not used because the curves did not terminate at the same point. The estimated number of RPA molecules was determined as follows. First, the peak intensity for an RPA binding event was calculated. Next, a global background signal was then subtracted from the intensity and then divided by the intensity of a single mCherry fluorophore, which has been calculated for this microscope. This yielded an estimated number of RPA molecules. The RPA binding footprint on ssDNA was then used to estimate the overall size of the strand separation.

### Multiple Sequence Alignment of Rad54

Representative Rad54L protein sequences were obtained from UniProt (62) and aligned in Jalview (2.11.3.2)(63) using MUSCLE with default settings (**Figure 1B**) (for complete alignment, see **Supplemental File 1**) (64). Rad54L from *Homo sapiens* (UniProt ID: Q92698) was also used to query the non-redundant database of Eukaryotic organisms (Taxonomic ID: 2759) using BLASTp (v. 2.15.0) (4), and the top 1,000 hits with an E-value less than 0.05 were aligned in Jalview (2.11.3.2)(65) using MUSCLE with default settings (**Supplemental File 1**). Only nine (9/1000) hits lacked one of the conserved residues (Arg, Tyr, Asp); eight are additional isoforms of hits containing the complete triad. The last sequence (EHH14742.1) lacks the Asp residue and is from *M. mulatta,* which encodes a second Rad54-like protein (UniProt ID: F7BLY5) that contains all three residues (**see Figure 1B**).

**Figure 1:**
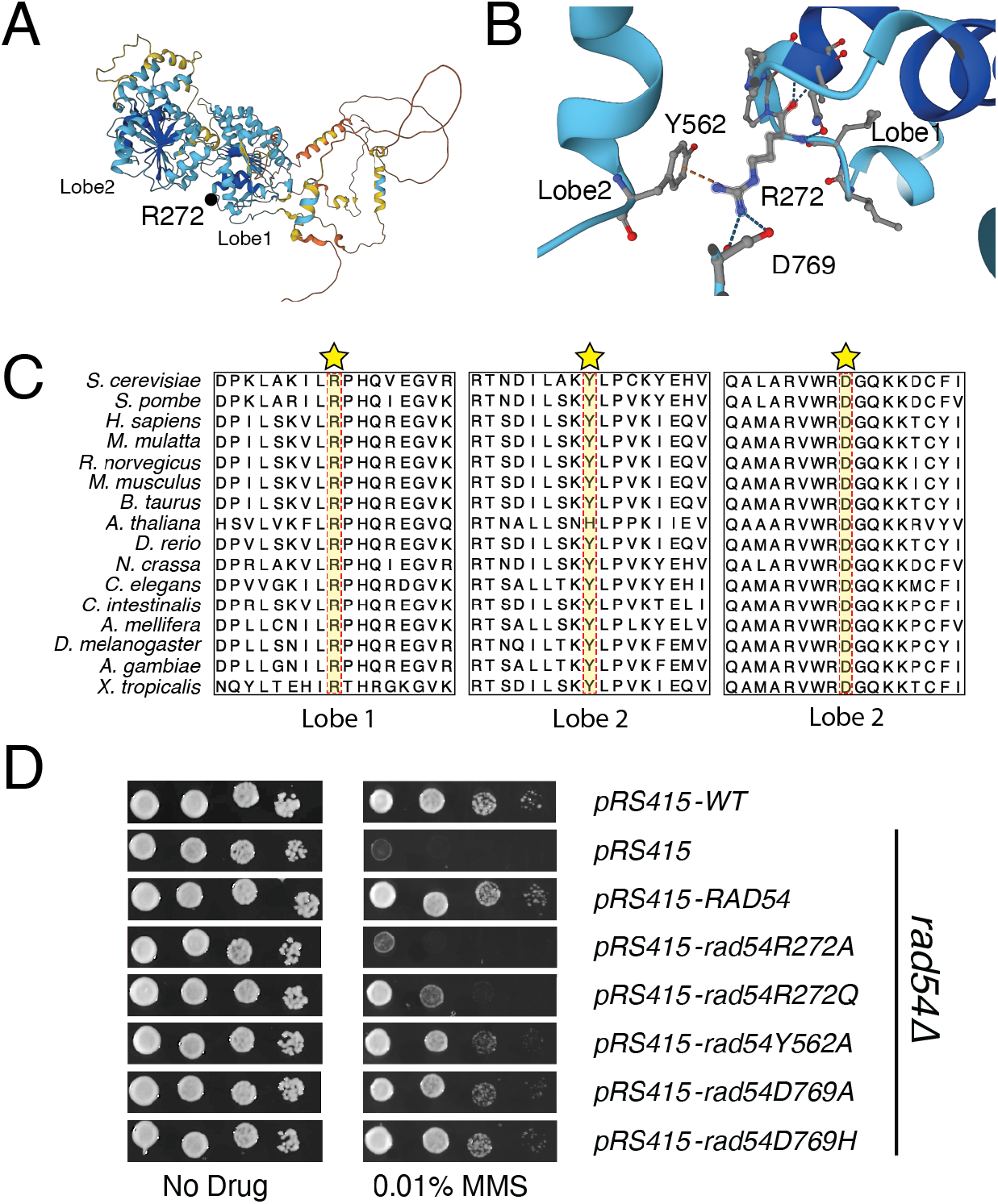
Disruption in the connection between Rad54 lobes leads to increased MMS sensitivity. **(A).** Alphafold2 structure of yeast Rad54. The protein is colored corresponding to the LDDT format. A black dot illustrates the site of mutation. **(B).** Expanded view within Rad54 that corresponds to a motif that bridges the two RecA lobes. The R272 and D769 residues were mutated in the COSMIC database. **(C).** Three regions of an amino acid sequence alignment of several eukaryotic Rad54 proteins showing the conservation of the R272, Y562, and D769 residues. Protein sequences were obtained from Uniprot and aligned in Jalview (2.11.3.2) using MUSCLE with default settings **(D).** Yeast complementation assay to evaluate Rad54 mutants for their ability to restore resistance to MMS. Alleles tested includes *RAD54*, *rad54R272A*, *rad54R272Q*, *rad54Y562A*, *rad54D769A*, and *rad54D769H*.

## Results

A primary goal of our study was to identify the reduction of function alleles in Rad54. To do this, we initially surveyed the COSMIC database (66). This is a repository for amino acid substitutions identified in human cancers. In selecting residues to mutate, we did not assign a cut-off for the number of cancers the amino acid substitutions were found in or define whether it was a driver or passenger mutation (**Supplemental Figure 1A**). We also confirmed that the identified residues were conserved across all eukaryotes. In total, we identified 60 amino acid substitutions in Rad54 that fit these criteria. We initially mutated thirteen residues in *S. cerevisiae RAD54* (**Supplemental Table 1, which includes equivalent human amino acid**) to test the complementation of sensitivity to methylmethanesulfonate (MMS) in *rad54*Δ strains. The four amino acid substitutions that failed to complement fully were *rad54P43S*, *rad54P43L*, *rad54R272Q*, and *rad54E381K* (**Supplemental Figure 1B**). While the substitution of proline at position 43 was interesting, we did not pursue this residue further because other mutations at this site were not sensitive to MMS. Likewise, the *rad54E381K* allele is a complete loss of function. Based on the *D. rerio* crystal structure, this mutant likely disrupts an essential alpha helix within one of the RecA lobes (**Supplemental Figure 2AB**). This left the *rad54R272Q* allele as a potential reduction of function allele that satisfied our criteria. Interestingly, this mutation has been identified in different malignant melanoma samples, and additional amino acid substitutions were identified in the COSMIC database. These included thyroid (R->L), as well as bladder, brain, and cervical cancer (R->W). We did not test these mutations directly, but the incidence of multiple variants at this site suggests a critical interaction (66).

The Rad54R272 residue is in the ATPase core of the Rad54 structure, and we were able to evaluate the position of this residue in the existing *D. rerio* structure (67) and an Alphafold2 generated structure of the *S. cerevisiae* Rad54 protein. From the structure, R272 formed a cross RecA lobe interaction with residues Y562 and D769 (**Figure 1AB**). Interestingly, the D769 residue was also identified in the COSMIC database as a D769H variant in breast cancer. We also evaluated the conservation of these three residues and found that all three were conserved in all eukaryotes, with an exception. In *Arabidopsis thaliana,* the tyrosine residue was changed to a histidine (**Figure 1C and Supplemental File 1**). This suggests that this tripartite interaction is conserved in eukaryotes and is likely fundamental to Rad54 biology and genome maintenance. We hypothesized that the R272Q mutation partially reduced function due to a loss of the charge interaction with the D769 mutation. However, this substitution could also potentially maintain a π-stacking interaction with the Y562 residue. If this were true, we would expect a further reduction of function if the R272 site was mutated to alanine.

We generated additional alleles including *rad54R272A, rad54Y562A, rad54D769H, and rad54D769A* (**Figure 1D**). We tested these alleles for MMS sensitivity and found that, as expected, the *rad54R272A* allele was more MMS sensitive than the *rad54R272Q* allele (**Figure 1D**). Likewise, we observed that the *rad54D769H*, *rad54D769A*, and *rad54Y562A* alleles had increased MMS sensitivity but were not as sensitive as the rad54R272Q and *rad54R272A* substitutions (**Figure 1D**). From this, we conclude that this interaction is required for fully functional Rad54, and we propose that this interaction is important for latching the two RecA lobes. We will refer to this interaction as the Rad54 latch motif. To further validate this hypothesis, we integrated the *rad54R272Q* and *rad54R272A* alleles into the genome and tested the complementation of the MMS sensitivity phenotype. We found that, as with the plasmid-based complementation, integration resulted in strains that were sensitive to MMS (**Supplemental Figure 3A**). From these results, we conclude that amino acid substitutions in this region of Rad54 result in a reduction of function.

Overexpression of Rad51 is partially toxic in *rad54*Δ cells. The mechanism behind this is the accumulation of Rad51 on chromosomes, which prevents proper chromosome segregation during cell division (42). We next tested whether the identified alleles of Rad54 were also sensitive to Rad51 overexpression by expressing the Rad51 protein from a galactose inducible promoter in *rad54R272Q* and *rad54R272A* strains. We found that the *rad54R272Q* and *rad54R272A* strains were able to complement the sensitivity caused by Rad51 overexpression observed in the *rad54*Δ strains (**Supplemental Figure 3B**). From this experiment, we conclude that these mutations in Rad54 can remove Rad51 from dsDNA without damage and are likely separation of function mutants.

### Mutations in the Rad54 latch reduce translocation efficiency on dsDNA

To test our hypothesis that these amino acid substitutions resulted in a reduced Rad54 translocation rate, we proceeded with biochemical characterization of the Rad54R272Q and Rad54R272A. The purified mutants bound to dsDNA as efficiently as WT (**Supplemental Figure 4A**) but exhibited reduced ATP hydrolysis activity (**Supplemental Figure 4BD)**. However, their ATPase was still stimulated by adding Rad51 (**Supplemental Figure 4CD**). This indicates that they are defective in hydrolysis but still interact with Rad51. We next used DNA curtains (60, 61) to characterize the WT-Rad54, Rad54R272Q, and Rad54R272A activity. This single molecule approach allows us to measure the DNA binding, speed, and translocation distance of Rad54 molecules (**Supplemental Figure 5A**). We initially characterized the ability of Rad54, Rad54R272Q, and Rad54R272A to bind dsDNA and found no defect in dsDNA binding for the Rad54 variants (**Supplemental Figure 5B**). We measured the translocation velocity of WT-Rad54 and found it to be 65+/−67 bp/sec with a mean track length of 3.7+/−2.5 kbp (**Supplemental Figure 5DE**). These values are consistent with previous observations (8). However, we could not measure Rad54R272Q or Rad54R272A as they did not move far enough to calculate an accurate velocity (**Supplemental Figure 5DE**). From these data, we conclude that Rad54R272Q and Rad54R272A have reduced translocation capacity.

We next evaluated the activity of Rad54R272Q and Rad54R272A in the context of Rad51. We have developed an *in vitro* single molecule assay to monitor Rad54 activity during the homology search and DNA alignment step of HR using DNA curtains (**Figure 2A**). In this experiment, Rad51 is preincubated with a 90 nt ssDNA oligo labeled with Atto647N. This allows the formation of short recombinase filaments or pre-synaptic complexes (PSCs). These structures are mixed with GFP-Rad54 and RPA-mCherry and flowed onto a preformed DNA curtain made from lambda dsDNA. This configuration allows direct observation of the Rad51-PSC bound by Rad54 as it searches for a homologous DNA sequence. In this experiment, we can measure the velocity and distance traveled by the Rad51/54-PSC. When using labeled RPA, we can also measure the association of RPA with the PSC, the time to RPA association, and the RPA binding lifetime. Finally, when a homologous sequence to the lambda DNA is used, we can also measure the probability of identifying the homologous sequences (**Figure 2B**). These measurements allow us to directly monitor the *in vitro* homology search.

**Figure 2:**
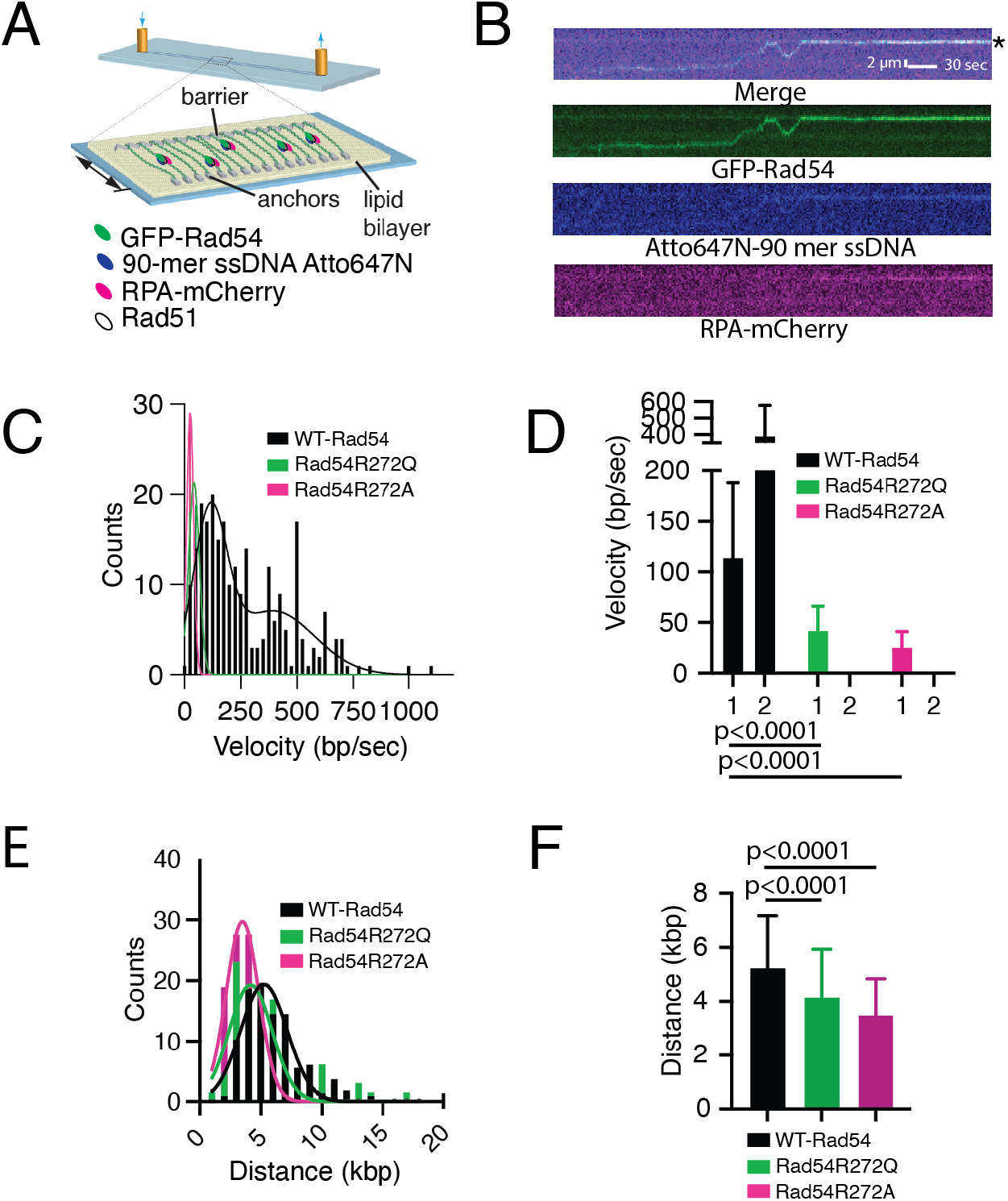
Disruption of the Rad54 latch reduces translocation velocity and distance. **(A).** Cartoon diagram illustrating DNA curtains experiment to monitor Rad51/Rad54 PSCs during the homology search step of HR **(B).** Representative Kymograph illustrates GFP-Rad54 (Green) as part of the Rad51-PSC translocation along DNA in search of homology. The recipient 90-mer ssDNA Atto647N (Middle) is labeled in Blue, and RPA-mCherry (Bottom) is included in the reaction (magenta). The Asterix denotes the sites of homology. **(C).** Distributions of measured velocities for WT-Rad54 (N=248), Rad54R272Q (N=65), and Rad54R272A (N=58). Gaussian distributions fit the data. **(D).** Bar graph representing the mean velocities calculated from a Gaussian distribution for Rad51/54-PSCs WT-Rad54 (N=248) (Black), Rad54R272Q (N=65) (Green), and Rad54R272A (N=58) (Magenta). 1 = the slower population of Rad54 observed in WT-Rad54 and 2 = the faster population observed in WT. **(E).** Distributions of measured translocation distances for WT-Rad54 (N=248), Rad54R272Q (N=65), and Rad54R272A (N=58). A Gaussian distribution fits the data. **(F).** Bar graph representing the mean distance traveled for Rad51/54-PSCs with WT-Rad54 (N=248), Rad54R272Q (N=65) (Green), and Rad54R272A (N=58) (Magenta). The error bars represent the standard deviation of the data.

For WT-Rad54, we measured the velocities of two unique populations. The faster of the two populations had a mean velocity of 394 +/− 183 bp/sec, and the slower one had a mean velocity of 114.3+/−73 bp/sec (**Figure 2CD**). Rad54R272Q and Rad54R272A had only a single population of velocities with a mean of 42.42+/−23.73 bp/sec and 25.7+/−15.32 bp/sec, respectively (**Figure 2CD**). We also measured the mean distance traveled by a unique Rad51/Rad54-PSC before stopping or dissociating. We found that the WT-Rad54 traveled 5.2+/−1.9 kbp. This value is consistent with previous reports (8). We observed a small but significant change in distance traveled with Rad54R272Q (4.1+/−1.7 kbp, p<0.0001) and Rad54R272A (3.4+/−1.3 kbp, p<0.0001) (**Figure 2EF**). From these data, we conclude that Rad54R272Q and Rad54R272A PSCs are defective in translocation on dsDNA.

### Rad54 mutants are impaired in donor DNA strand opening

It has previously been shown that as Rad51/54 PSCs move along DNA, they can recruit RPA. This is not through protein-protein interaction and likely results from the generation of locally underwound DNA behind translocation and overwound DNA ahead of translocation (8, 15) (**Figure 3A**). We evaluated how Rad54R272Q and Rad54R272A affected RPA dynamics during PSC translocation (**Figure 3B**). We estimated the number of RPA-mCherry molecules associated with translocating PSCs by comparing their overall intensities with known intensities from individual photobleaching experiments (68). This analysis measured the mean number of RPA molecules bound to PSCs at 2.5+/−1.5 (**Figure 3C**). This value was like previously reported values (8). In contrast, Rad54R272Q and Rad54R272A had RPA values of 2.13+/−1.2 and 1.75+/−1.3, respectively (**Figure 3C**). These values represent a significant difference from WT. If we quantize the amount of RPA, then WT-Rad54 will likely have 2-3 molecules of RPA bound, representing underwound DNA of 60-90 nt. In contrast, the mutant versions of Rad54 are likely to have 1-2 molecules of RPA, suggesting an underwound region of 30-60 nt. From these data, we conclude that the Rad54 substitutions have reduced strand opening activity.

**Figure 3:**
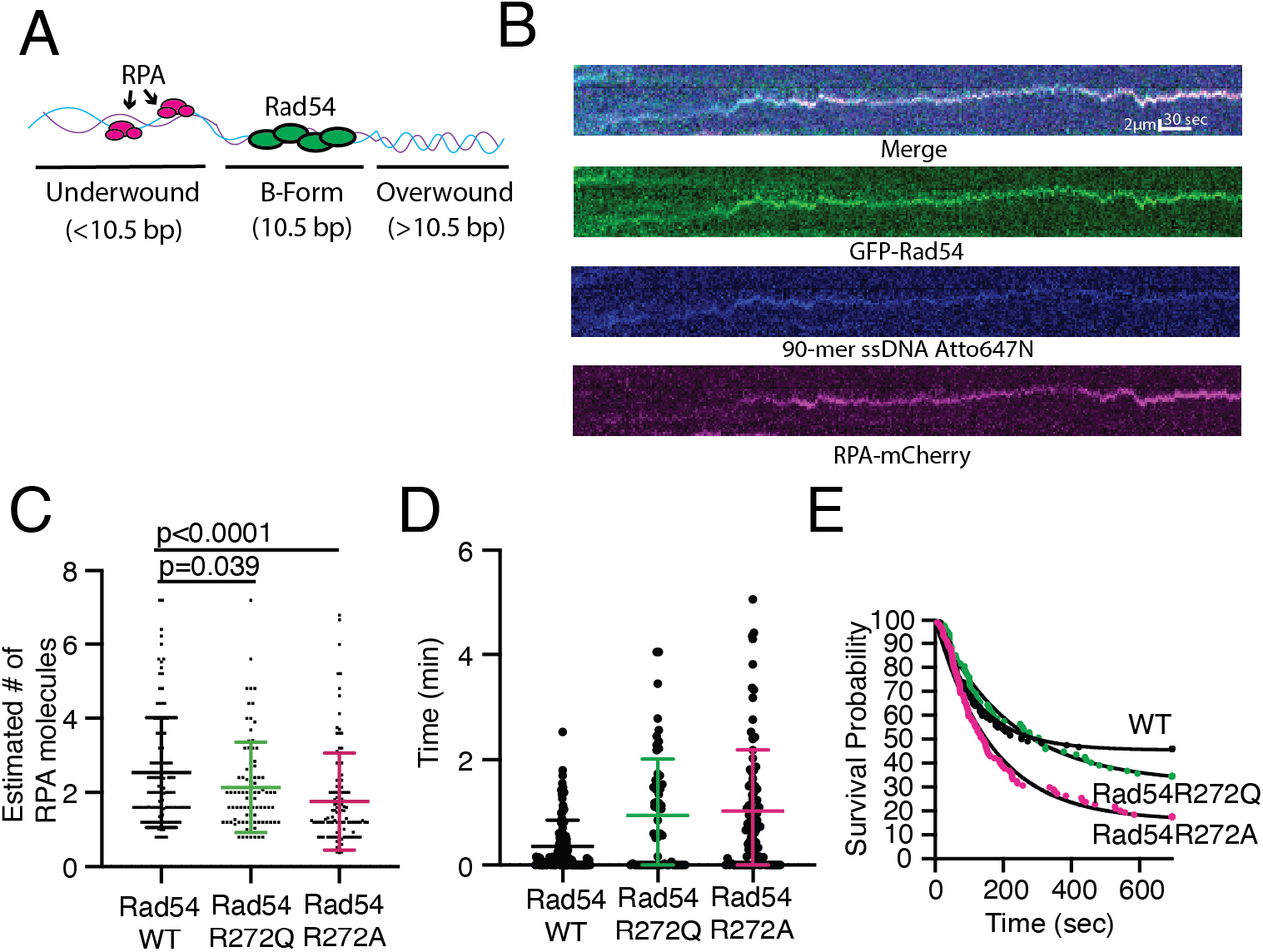
Mutations in Rad54 influence RPA dynamics on the PSC. **(A).** Cartoon schematic illustrating the proposed model for Rad54 movement on dsDNA. Movement creates underwound DNA behind and overwound DNA ahead. **(B).** Representative kymograph illustrating co-translocation of GFP-Rad54 (Green), 90-mer ssDNA-Atto647N (Blue), and RPA-mCherry (Magenta). **(C).** Dot plot representing the estimated number of RPA molecules based on photobleaching measurements for an individual step size for WT-Rad54 (N=119), Rad54R272Q (N=87) (Green), and Rad54R272A (N=114) (Magenta). The cross and error bars represent the mean and standard deviations of the data. **(D).** Dot plot representing the time it takes for RPA to associate with Rad51/Rad54-PSCs that are bound to dsDNA for WT-Rad54 (N=108) (Black), Rad54R272Q (N=60) (Green), Rad54R272A (N=95) (Magenta). The cross and error bars represent the mean and standard deviation of the data. **(E).** The survival probability distribution for the dissociation rate of RPA from WT-Rad54 (N=122) (Black), Rad54R272Q (N=87) (Green), and Rad54R272A (N=114) (Magenta).

We next evaluated the kinetics of RPA association with the PSC. We measured the time it took to observe RPA associating with the PSC after it bound to the dsDNA. We found that the median association time for WT was 0.14 min (**Figure 3D**). In contrast, the Rad54R272Q and Rad54R272A mutants had median association values of 0.6 and 0.73 min, respectively (**Figure 3D**). These data likely reflect that these Rad54 mutants are delayed in their ability to generate underwound DNA. We also measured the percentage of RPA molecules that dissociated after binding. For WT Rad54, we found that 55% of RPA molecules dissociated after binding (**Figure 3F**). In contrast, 69% of RPA molecules dissociated after binding in the case of the Rad54R272Q mutant, and 84% of RPA molecules dissociated in the case of the Rad54R272A mutant (**Figure 3F**). Together, these data suggest that these Rad54 substitutions have a reduced ability to generate and stabilize strand-separated DNA.

### A reduction in motor function alters DNA sequence alignment

Identification of the correct DNA sequence during the homology search can occur through 1D or 3D movement of the PSC (8, 10). In our assay, we can identify successful searches that have occurred through 1D movement by determining if there is movement along the DNA before sequence recognition (**Figure 4A**). We also observe sequence recognition events that occur instantaneously (**Figure 4A**). We interpret these recognition events to mean the sequence has been identified by 3D movement. We cannot exclusively rule out a short/fast translocation event faster than the resolution of our experiment. However, this interpretation is consistent with defects observed in the Rad54 mutations.

**Figure 4:**
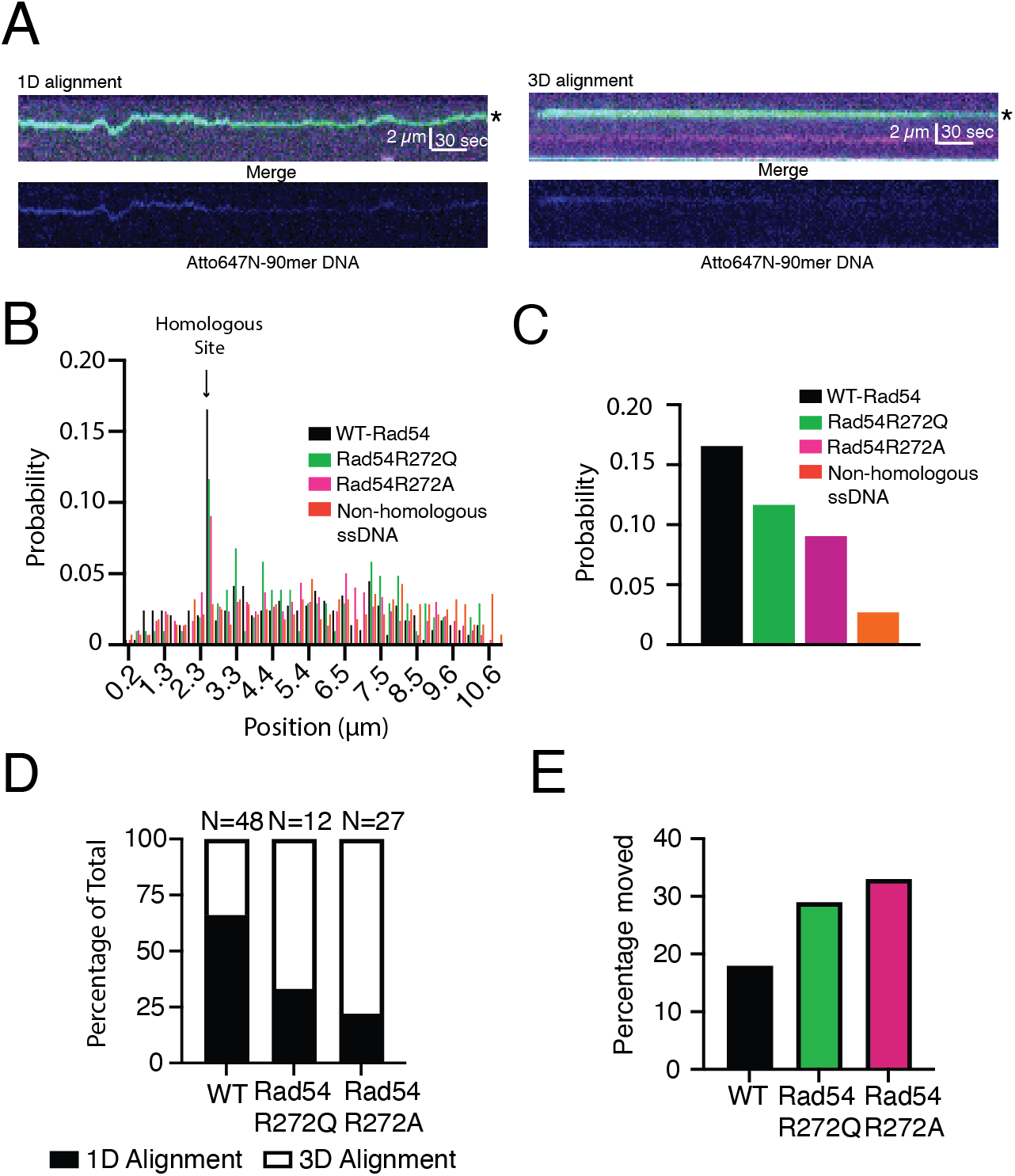
Reduced Rad54 activity alters the efficiency of DNA sequence alignment. **(A).** Representative kymographs illustrating the alignment of the homologous sequence by 1D movement (Left) and 3D movement (Right). The kymographs represent a merged three-color image (Top) of GFP-Rad54 (Green), RPA-mCherry (Magenta), and 90-mer ssDNA Atto647N (Blue) and an individual image of the Atto647N-90-mer (Bottom). A star denotes the site of homology. **(B).** Graph representing the binding probability distribution of PSCs across lambda DNA after 15 minutes. An arrow marks the site of homology. The data are for Rad51/54-PSCs consisting of WT-Rad54 (N=290) (Black), Rad54R272Q (N=103) (Green), and Rad54R272A (N=298) (Magenta). Non-homologous sequence (N=280) (Orange) **(C).** The expanded view of the graph in B represents only the homologous site. **(D).** Graph representing the percentage of sequence alignment events that occur via 1D movement versus 3D movement for WT-Rad54, Rad54R272Q, and Rad54R272A. **(E).** Bar graph illustrating the number of PSCs that move after dwelling on the homologous site for 10 seconds.

From these observations, we can also determine the efficiency of DNA alignment through both 1D and 3D movement. We first calculated the global frequency of DNA sequence alignment by monitoring the Rad51/54-PSC dwelling at the correct region of homology versus all possible binding sites after fifteen minutes of observation. These values are compared to a non-homologous piece of DNA to determine whether the predicted site is enriched. In these experiments, the WT-Rad54 is aligned with the homologous site with a probability of 0.165. This is 5.75-fold higher than the measured probability of 0.0285 for non-homologous DNA and 7.5-fold higher than the mean probability for all non-homologous sites,0.022 (**Figure 4BC and Supplemental Figure 6ABCD**). In contrast, the Rad54R272Q and Rad54R272A had a probability of 0.11 and 0.09, respectively. This represented a 1.5-fold and 1.8-fold reduction from WT-Rad54, respectively (**Figure 4BC and Supplemental Figure 6ABCD**). From these data, we conclude that the Rad54 mutations impair DNA sequence alignment, likely due to a reduction in motor function.

We next evaluated the population of molecules at the target site that aligned via 1D movement versus 3D movement. For WT-Rad54, 66% (32/48) were aligned by 1D movement, and 34% (16/48) were aligned by 3D movement (**Figure 4D**). By comparison, Rad54R272Q aligned DNA by 1D movement 33% (4/12), and Rad54R272A aligned DNA by 1D movement 22% (6/27) of the time. These data suggest that DNA sequence alignment is reduced in these Rad54 mutations and that there is a change in the mechanism used to align DNA. We also measured the number of molecules that came into alignment at any point during the experiment for greater than twenty seconds but then dissociated. In the case of WT-Rad54, this number was 18% (11/59) (**Figure 4E**). For Rad54R272Q and Rad54R272A, it was 29% (5/17) and 33% (13/40), respectively. From these data, we conclude that a reduction in the rate of Rad54 translocation leads to a diminished capacity to align DNA stably.

### Rad54 mutants require longer stretches of homology to form stable D-loops

Our single molecule analysis indicates that Rad54R272Q and Rad54R272A have reduced capacity to translocate on DNA, align DNA, and stabilize strand opening events. These data suggest that these proteins should be deficient in D-loop formation. To test this question, we used an *in vitro* D-loop formation assay (**Figure 5A**). This assay is based on the pairing and formation of short D-loops using the pUC19 plasmid as a substrate. We performed the experiments with 65, 90, and 130-nt of homology. Each homologous ssDNA contained a 25 nt non-homologous stretch at the 5’ end of the DNA that was annealed with a 25 nt oligo labeled with Atto647N (**Figure 5A**).

**Figure 5:**
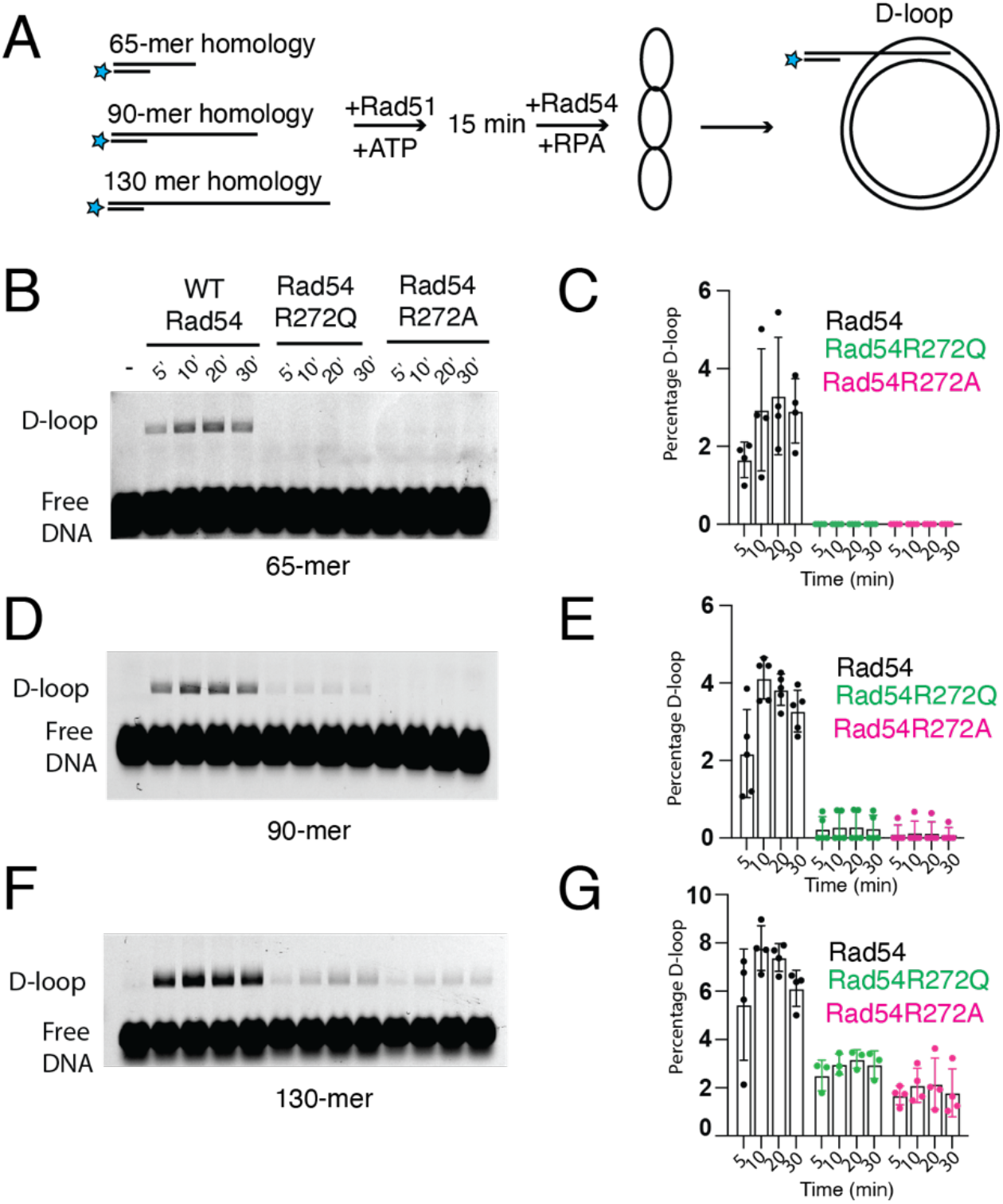
Rad54 mutants have a diminished ability to form D-loops *in vitro*. **(A).** Cartoon diagram illustrating experiment to measure WT-Rad54, Rad54R272Q, and Rad54R272A activity for D-loop formation with different lengths of homology. **(B).** Representative gel illustrating D-loop formation for WT-Rad54, Rad54R272Q, and Rad54R272A with a 65-mer of homology to the pUC19 plasmid. **(C).** The bar graph quantifies the D-loop formation percentage for WT-Rad54, Rad54R272Q, and Rad54R272A. The error bars represent the standard deviation of 4 independent experiments. **(D).** Representative gel illustrating D-loop formation for WT-Rad54, Rad54R272Q, and Rad54R272A with a 90-mer of homology to the pUC19 plasmid. **(E).** The bar graph quantifies the percentage of D-loop formation for WT-Rad54, Rad54R272Q, and Rad54R272A. The error bars represent the standard deviation of 5 independent experiments. **(F).** Representative gel illustrating D-loop formation for WT-Rad54, Rad54R272Q, and Rad54R272A with a 130-mer of homology to the pUC19 plasmid. **(G).** The bar graph quantifies the D-loop formation percentage for WT-Rad54, Rad54R272Q, and Rad54R272A. The error bars represent the standard deviation of 3-4 independent experiments.

We first measured the 65-mer homologous sequence and found that WT could promote stable D-loop formation. In contrast, we failed to observe measurable D-loop formation with Rad54R272Q and Rad54R272A (**Figure 5BC**). We next measured 90 nt of sequence homology. As with the 65-mer, the WT-Rad54 promoted the formation of stable D-loops. For RadR272Q in some experiments (n=2/5), we observed stable D-loop formation with a 5-fold reduced efficiency relative to WT-Rad54 (**Figure 5DE**). For the Rad54R272A mutant, we observed measurable D-loop formation in fewer (n=1/5) experiments, and in this experiment, there was a 6-fold reduction relative to WT-Rad54. When using a 130 nt homology, we were able to see consistent D-loop formation for both the Rad54R272Q (n=3/3) and the Rad54R272A mutations (n=4/4) (**Figure 5FG**). However, both mutants formed D-loops with reduced efficiency, 2.3-fold for Rad54R272Q and 3.5-fold for Rad54R272A, relative to WT-Rad54. From these experiments, we conclude that the Rad54R272Q and Rad54R272A proteins have reduced capacity to form stable D-loops and that longer tracts of homology can begin to alleviate the reduction of this activity. This is consistent with the role of Rad54 in stabilizing shorter early pairing events.

### Reduction in Rad54 function leads to increases in crossover outcomes

The HR reaction proceeds through a series of intermediates determining whether DNA sequence information is passed between donor and recipient DNA molecules. These intermediates primarily depend on the capture of the second DNA end. For example, if the D-loop is disrupted, the recipient DNA is annealed to the second broken end and used as a repair template as part of the SDSA pathway. This results in non-crossover (NCO) outcomes. If the D-loop is not disrupted and the non-template strand captures the second end, a double Holliday junction (dHJ) can form. This structure can result in crossover (CO) or NCO outcomes. If the second end is not located or does not exist, break induced reapplication (BIR) can occur, a mutagenic long-range replication event that can result in loss-of-heterozygosity outcomes.

Rad54 is believed to promote NCO outcomes as part of the synthesis-dependent strand annealing (SDSA), which protects genomic integrity by preventing excessive sequence exchange between donor and recipient DNA molecules. Previous reports have shown that overexpression of *RAD54K341R* has a dominant negative effect on WT strain, causing increases in CO outcomes in diploid yeast (46). This could be due to the disruption of Rad54 ATPase activity, as this motor functions as a multimer (8, 69). We hypothesized that *rad54R272Q* and *rad54R272A* mutants may have similar phenotypes and could result in higher CO levels.

We used a genetic reporter assay that monitors outcomes of allelic exchange between homologous chromosomes (27, 70). Strains used in this assay have ade2 (ade2-I and ade2-n) heteroalleles located on chromosome XV. One of these alleles has an I-Sce1 cleavage site (ade2-I), and the other has a disrupted reading frame toward the start of the gene (ade2-n). Double strand breaks can be induced by the expression of the I-Sce1 nuclease under the control of a galactose inducible promoter. In this experiment, both sister chromatids are cut, which results in the ade2-n gene on the homologous chromosome being used as a repair template. This results in gene conversion events. Short tract gene conversion events, <1 kbp, result in recovery of an active *ADE2* gene and white colonies. Long tract gene conversion outcomes, >1 kbp, result in the conversion of the ade2-I to ade2-n, which maintains an inactive copy of the gene. Each sister chromatid can undergo gene conversion through either long or short-tract repair. If each is repaired by the same type, then the colony is either red or white. However, if one sister is fixed by short tract and the other by long, the colonies can be red/white sectored. These colonies are most important for inferring recombination outcomes because they have undergone division before plating.

Each homologous chromosome contains a different antibiotic marker to determine the outcome of a recombination reaction. Non-crossover (NCO) outcomes occur when there is no change in the segregation pattern of the antibiotic markers because the chromosomes remain heterozygous. Crossover (CO) outcomes can be inferred when each part of a sectored colony is sensitive to different antibiotics. This occurs because antibiotic markers have switched chromosomes, and chromosome VX in this colony is now homozygous for one antibiotic marker. This loss-of-heterozygosity (LOH) event can occur with long-track or short-track repair. Another outcome that can be determined from this assay is break-induced replication (BIR). This can be inferred in a sectored colony when a red sector is resistant to both antibiotics, and the white sector is only resistant to one. This represents a LOH event in which a long-track repair event has copied the antibiotic marker during repair (**Figure 6AB**) (71). This assay also contains markers to report on chromosome loss. We have previously used this assay and found that *rad54*Δ*/rad54*Δstrains are severely defective in this assay and result in limited recombination (27).

We compared the *rad54R272Q* and *rad54R272A* alleles to the WT for their ability to promote recombination (**Supplemental Figure 7A and Supplemental Table 4-6**). In contrast to the *rad54*Δ these strains promoted viable recombination outcomes. For sectored recombinants, WT resulted in 50.4 +/−2.7% CO (**Figure 6C**). In contrast, the *rad54R272Q* substitution resulted in 61+/−2.8% CO, and the *rad54R272A* substitution resulted in 64+/−4.5% CO (**Figure 6C and Supplemental Table 4-6**). The *rad54R272A* allele had slightly increased chromosome loss, which could account for the difference from the *rad54R272Q* allele (**Supplemental Table 4-6**). For solid red recombinants, WT strains only resulted in CO outcomes 13.2+/−4.6 % of the time. The *rad54R272Q* and *rad54R272A* alleles displayed increased CO outcomes with 30.3+/−10 % and 28.3+/− 9%, respectively (**Figure 6D and Supplemental Table 4-6**). When we combined the total recombinant population, we observed that 40+/−1.1% of recombinants resulted in CO outcomes in WT. In the rad54R272Q and rad54R272A mutants these numbers were 50.4+/−4.0 % (p=0.0003) and 56.3+/−3.4 % (p<0.0001) respectively (**Figure 6E**). Together, these data suggest that reducing Rad54 activity increases CO outcomes. We also measured BIR outcomes and observed a 2-fold reduction in the BIR observed in the *rad54R272Q* and *rad54R272A* relative to WT (**Supplemental Figure 7BCD**). This is consistent with previous reports that the rate of strand invasion may affect BIR outcomes, and these data suggested that *rad54R272Q* and *rad54R272A* may promote strand invasion at reduced rates.

**Figure 6:**
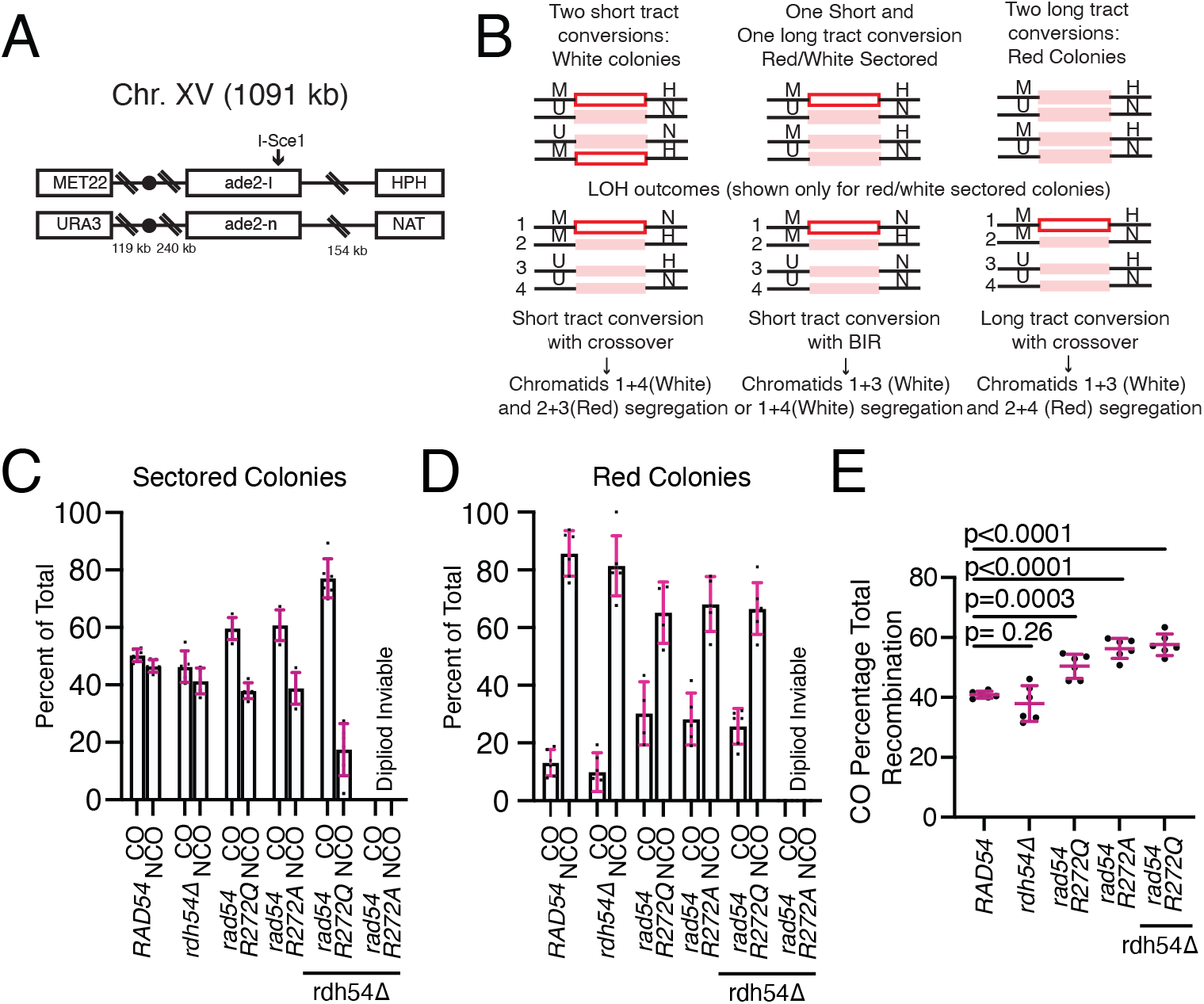
Mutations in Rad54 lead to increased crossover outcomes. **(A).** Schematic diagram illustrating the double strand break reporter construct used in these experiments. **(B).** Schematic diagram illustrating the possible recombination outcomes using this assay. The general outcome from recombination can be inferred from the color of the colony and the antibiotic sensitivity. **(C).** Graph illustrating the CO and NCO outcomes for sectored colonies of WT, *rdh54Δ, rad54R272Q*, *rad54R272A*, *rad54R272Q rdh54Δ,* and *rad54R272A rdh54Δ.* The error bars represent the standard deviation of six independent experiments. **(D).** Graph illustrating the CO and NCO outcomes ratio for solid red colonies of WT, *rad54R272Q, rad54R272A, rdh54Δ, rad54R272Q rdh54Δ, and rad54R272A rdh54Δ.* The error bars represent the standard deviation of six independent experiments. **(E).** Graph illustrating crossover outcomes as a percentage of total recombination events for WT, *rad54R272Q, rad54R272A, rdh54Δ, rad54R272Q rdh54Δ, and rad54R272A rdh54Δ.* The bars represent the mean, and the error bars represent the standard deviation from six independent experiments.

Rdh54 is a paralog of Rad54 and has previously been implicated in controlling the length of strand invasion reactions by regulating Rad54 during D-loop formation (14). We hypothesized that deletion of *RDH54* would suppress the *rad54R272Q/A* phenotypes, and this may allow longer, more stable invasion products to form. As has previously been reported, there was little impact on changes in CO in the *rdh54*Δ strains (**Figure 6CD**). We also observed a significant increase in BIR outcomes (**Figure 6E**). We were surprised that when we tested the *rad54R272Q rdh54*Δ double mutant, there was a slight increase in total CO events to 57.6+/−3.6. This was larger than the *rad54R272Q* strain (**Figure 6CDE**). This was inconsistent with our hypothesis but could be explained if Rdh54 had other impacts on the recombination reaction. For example, the deletion of this gene may further inhibit the SDSA pathway (13, 14) or lead to an excess of uncontrolled Rad51 molecules, creating longer Rad51 filaments on ssDNA (41, 72). We did observe a restoration of BIR outcomes to the WT level (**Supplemental Figure 7BCD**), which is consistent with the initial hypothesis. Unfortunately, after several attempts, we could not get healthy diploids with the *rad54R272A rdh54*Δ double mutants. This is consistent with previous reports that *rad54Δrdh54*Δ has a reduction in diploid growth. These observations are consistent with a strain that has become too defective to support viability in the diploid state. From these data, we conclude that a reduction in Rad54 activity results in increased genomic re-arrangements in the form of CO outcomes.

## Discussion

Here, we have identified and characterized the activity of Rad54 reduction of function substitutions that slow down the translocation activity of Rad54. This amino acid site was found in the COSMIC database from multiple cases of malignant melanomas. We further found that this residue is central to the Rad54 latch domain, which is conserved in eukaryotes. Substitutions at this site diminished Rad54 activity by reducing DNA sequence alignment by 1D movement and caused the formation of unstable abortive strand invasion intermediates. This leads to an increase in genomic rearrangements as characterized by CO outcomes. While our study pertains to Rad54 activity, it suggests that factors that promote kinetic regulation of early D-loop intermediates can balance HR outcomes.

### Cancer substitutions

We used the COSMIC database to identify amino acid substitutions in Rad54 that were reduction in function. What we discovered was a near-universally conserved three amino acid motif that caused a decrease in Rad54 translocation activity. This latch structure was located on the interface between the two RecA lobes of Rad54, with the R272 residue acting as the lynchpin to tie the lobes together. We hypothesize that this interaction is a charge-charge interaction between R272 and D769 and a π-stacking interaction between Y562 and R272. In the sole example of Rad54, that had a substitution in the latch, *A. thaliana* had a histidine where the tyrosine is found in all other species. We suspect this histidine can still form a stacking interaction with the arginine, conserving this part of the interaction. The D769 residue was also found in the COSMIC database mutated to a histidine.

Mechanically, Rad54 is intrinsically flexible and, like other Snf2 motors, can exist in an open and closed state (73). When bound to dsDNA and ATP, the lobes align, and the motor can translocate along dsDNA. Based on the location of the latch, these residues likely work together to prevent the two RecA lobes from drifting apart as the enzyme resets for the next round of hydrolysis and translocation (74). Disruption of the latch results in defective Rad54 translocation and reduced ATP hydrolysis. The universal conservation of the latch structure and the existence of mutations in human cancers indicate an interaction critical for genome maintenance in all eukaryotes. The consequence of losing this interaction is an increase in genomic rearrangements. While we cannot identify these amino acid substitutions as the cause of malignant melanoma in these patients, it would almost certainly have been an underlying factor in disease progression. While this is a specific case, it suggests that human cancers with low-functioning Rad54 may be prone to genomic rearrangements, as discussed below. Ultimately, while this may limit Rad54’s use as a potential drug target, these results offer new insight into its use as a potential diagnostic marker to evaluate the likelihood of genomic rearrangements for specific cancer types.

### 1D Translocation based homology search

The homology search occurs through multiple biophysical mechanisms, including 1D/3D diffusion and 1D translocation-based searches (8–10, 18, 75). The efficiency of 3D diffusive-based mechanisms will scale proportionally with the length of homology and concentration of non-specific competitor DNAs (9). Longer pieces of homology increase the probability of productive contact between the donor and recipient DNA. Rad54 can contribute to the homology search by increasing association with donor DNA (76) and providing a 1D translocation-based search that enhances sequence alignment efficiency (8).

1D translocation along dsDNA is severely compromised in the Rad54R272Q and Rad54R272A mutants. This causes reduced DNA sequence alignment by the Rad51/54-PSC and a diminished ability to form D-loops in bulk. Rad54 latch substitutions failed to create stable D-loop products on short homologous DNA sequences. However, as the size of the homologous sequence increased, we were able to observe the formation of stable D-loops. We interpret this result to mean that 1D translocation is important for sequence alignment *in vitro* and not the process of D-loop formation. If translocation were required to stabilize D-loop formation, we would have expected the opposite result, and the Rad54R272Q/A substitutions would have formed D-loops on shorter substrates instead of longer. This suggests that the strand-opening activity of Rad54 is critical for D-loop formation (discussed below), which can still function, with reduced efficiency, in the mutated proteins if there are 3D collisions between the recipient and donor DNA. However, regardless of the length of homology, translocation catalyzed the formation of D-loops by improving the homology search.

While the direct contribution of Rad54 translocation to the homology search remains to be defined *in vivo*, the increased efficiency of target selection afforded by 1D translocation may increase repair when the sister chromatid is the preferred template, increasing the kinetics of sequence alignment in a local context. Here, we have identified mutations in Rad54 that may contribute to decreased 1D homology search in cells, leading to sensitivity to DNA-damaging agents. Further work will be needed to nail down the precise *in vivo* role of Rad54 during the homology search.

### Rad54 can use topology to regulate sequence recognition probability

The superhelical density of donor dsDNA can regulate strand invasion and the general HR reaction (77, 78). Negative supercoiling of DNA causes strand underwinding characterized as <10.4 bp per turn of the double helix. The underwinding of DNA improves the activity of Rad51-mediated invasion by reducing the energy required to separate DNA strands and make each subsequent triplet pairing event between the donor and recipient DNA. Rad54 can regulate superhelical density by adding twist to DNA, likely generating negative supercoils behind translocation and positive supercoils ahead (15). This activity leads to local DNA melting, as observed by the association of RPA with moving Rad51/54 PSCs (8). Here, we have identified variants in Rad54 that generate shorter and less stable regions of underwound DNA. These lack efficient D-loop formation and are sensitive to the DNA-damaging agent MMS.

Current structural models show that RecA promotes progressive base pairing during strand invasion by separating the template and non-template strands of DNA through sequestration of the non-template strand by Loop 2. This allows additional homology pairing and extends the initial D-loop (79, 80). This model predicts that strand pairing occurs over a short distance with the probability of invasion termination occurring once every 15 bp (79, 80). This is consistent with observations made on hRAD51 in the post-synaptic state (81), and the Rad51 mechanism is likely similar. These short D-loops may be sufficient to promote repair in bacteria or may be stabilized by the cooperation of multiple short invasion products that accumulate over time. However, in eukaryotes, they may be kinetically unstable in the competitive environment of the nucleus (9). In this case, these pairing events may result in abortive invasion events that fail to extend and form stable D-loop intermediates competent for repair.

We propose a model where Rad54 uses ATP hydrolysis to promote topological changes in donor DNA that reduce abortive Rad51 invasions. In this case, strand separation promoted by Rad54 activity will catalyze the expansion of Rad51 invasion products and promote stabilization through further base pairing (**Figure 7AB**). The mechanism of catalysis is the generation of underwound DNA, decreasing the probability of termination during each base pairing event. Overall, this mechanism would improve the rate of the homology search by ensuring that early homology recognition events are converted to stable D-loop intermediates that are competent to promote repair. In the case of the Rad54 variants identified here, shorter, less stable regions of underwound DNA are formed. This likely lowers the probability of D-loop expansion and increases the time to stable product formation. We propose that reducing Rad54 biochemical activity increases abortive invasion, ultimately slowing the HR reaction rate (**Figure 7AB**). As discussed below, increases in abortive invasion may cause higher crossover outcomes.

**Figure 7:**
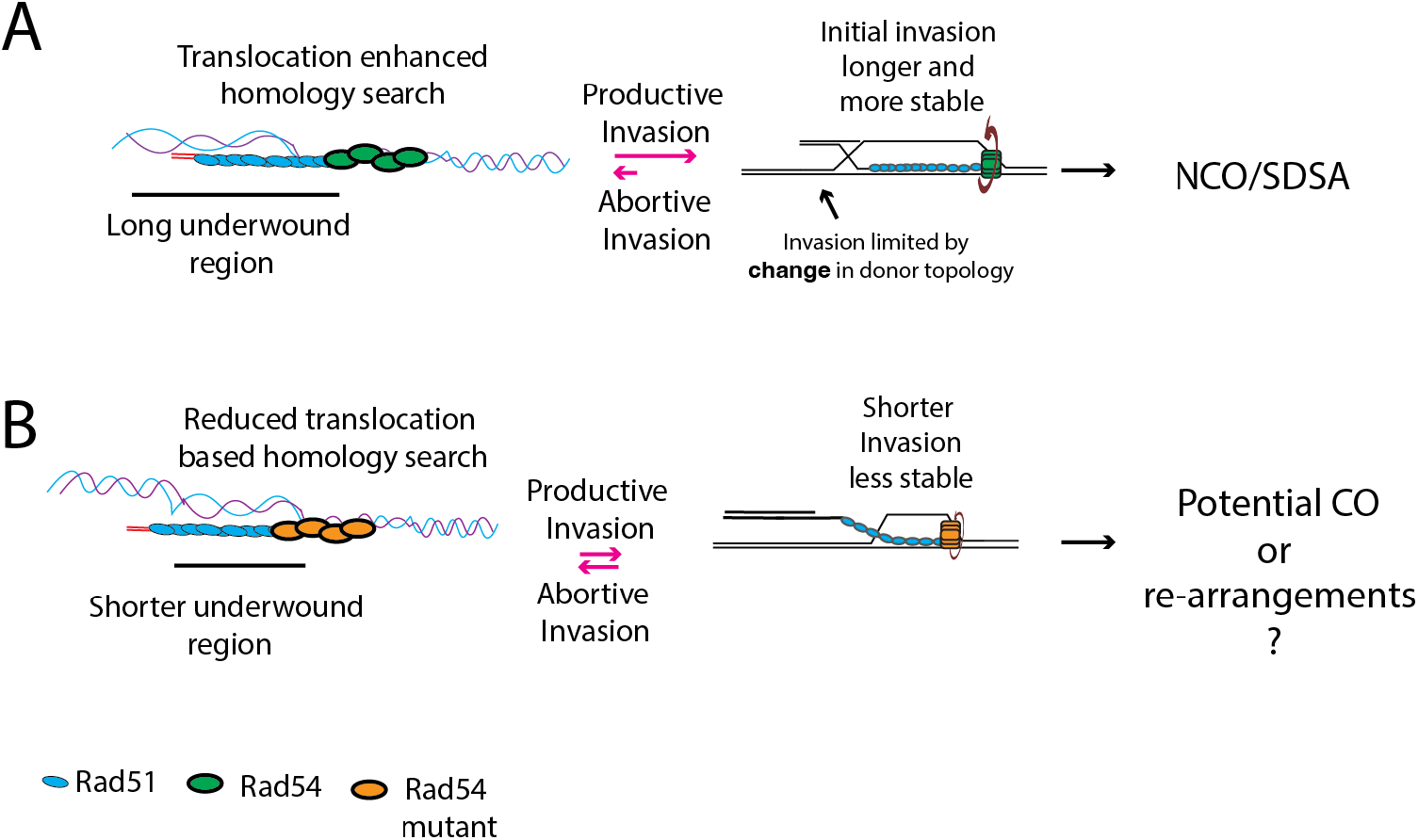
Model for reduction in abortive invasion. **(A).** Schematic diagram illustrating that longer tracts of underwound DNA generated by Rad54 lead to faster invasions and non-crossover repair or SDSA. **(B).** Mutant versions of Rad54 lack general translocation activity and produce shorter regions of underwound DNA, leading to less stable invasion products.

### Rad54 may serve as an anti-recombination factor

DNA motors that prevent extensive recombination between donor and recipient molecules are characterized as anti-recombinase enzymes. This includes Sgs1/Bloom helicase (BLM), Mph1 (FANCM), and the helicase Srs2 in *S. cerevisiae* (35, 82, 83). Deleting these genes leads to elevated CO outcomes between donor and recipient DNA. Rad54 is proposed as part of the SDSA pathway (84) through its branch migration activity. A two-step mechanism exists for Rad54 function at D-loops, with translocation aiding in both D-loop formation and disruption. We offer an alternate single-step model where Rad54 may create a window in the DNA, which catalyzes the strand invasion process, maintaining DNA distortion while Rad51 facilitates invasion (**Figure 7AB**). The window could be terminated at the interface between underwound DNA and B-form DNA outside the influence of Rad54 activity. This return to B-form DNA would make termination of invasion more likely. Loss of this window may promote an increased likelihood of CO formation.

We cannot specifically conclude the biological mechanism by which the reduction in the function of Rad54 leads to an increase in CO outcomes. One possibility is a kinetic mechanism in which a reduction in the rate of D-loop formation leads to prolonged HR and an increased probability of CO formation. Second, reducing Rad54 activity may increase the probability of double Holliday junction (dHJ) formation and, by extension, an increase in CO formation. Finally, we must consider that additional Rad54 may be recruited to the stable D-loop later in the reaction to promote the resolution of the D-loop. Further work will be needed to understand the specific genetic mechanism better.

Deletion of the *RAD54* paralog *RDH54* in the *rad54R272Q* background resulted in higher levels of CO. We initially thought that *RDH54* may serve as a suppressor of this phenotype by restoring the extended strand invasion tract length in the defective Rad54 alleles. Instead, we observed increased CO and further loss of NCO events. Rdh54 has been shown to stabilize early strand invasion intermediates and to reduce Rad54 mediated invasion tracts (14). This may be part of a role Rdh54 has in the SDSA pathway. Deletion of Rdh54 may further impair this pathway in the *rad54R272Q* background, suggesting these motors may cooperate to promote SDSA. Whether this is due to a direct physical interaction with the Rad51 filament, changes in global regulation of Rad51 pools, or regulation of a strand invasion window is unclear (14).

### Controlling the length of nascent D-loops is likely an essential feature of limiting genetic exchange

In *S. cerevisiae*, Rad54 is critical in developing proper Rad51 strand invasion intermediates. This co-dependence is less evident in higher eukaryotes, and additional factors may regulate the productive versus abortive invasion in higher eukaryotes. Factors that regulate early strand invasion intermediates include RAD51AP1 (85) and HOP2/MND1 (86, 87). These factors will likely regulate D-loop intermediates through distinct mechanisms. RAD51 in higher eukaryotes may also be more stable during initial strand invasion and more readily prevent abortive invasion events. Despite this, the amino acid substitution characterized here was identified in human cancer cases and is universally conserved. This suggests that Rad54 substitutions that increase abortive invasion may also contribute to genomic rearrangements in human cancers in pathways that utilize Rad54 to mediate repair.

## Data availability

All data is available upon request.

## Acknowledgments

We would also like to acknowledge David Moraga, Eric Alani, and Marcus Smolka for their critical manuscript reading. We would also like to thank members of the Cornell R3 group and the members of the Crickard laboratory for their helpful input during the project development.

## Author contributions

KS performed the initial screen, genetically characterized Rad54 mutants and MVW purified proteins, performed experiments, analyzed data, and helped write the manuscript. JH performed experiments and analyzed data. BF helped with the writing and editing of the manuscript. JS provided cloning support and critical strains. JBC performed experiments, analyzed data, provided reagents, and wrote the manuscript with input from KS, MVW, and JH. This work was supported by NIGMS R35142457 to JBC and Cornell Startup funds to JBC.

## Notes

### Competing Interest Statement

The authors have declared no competing interest.

