## Supplemental_Data for "The translocation activity of Rad54 reduces crossover outcomes during homologous recombination"

### Supplemental Figure 1

A

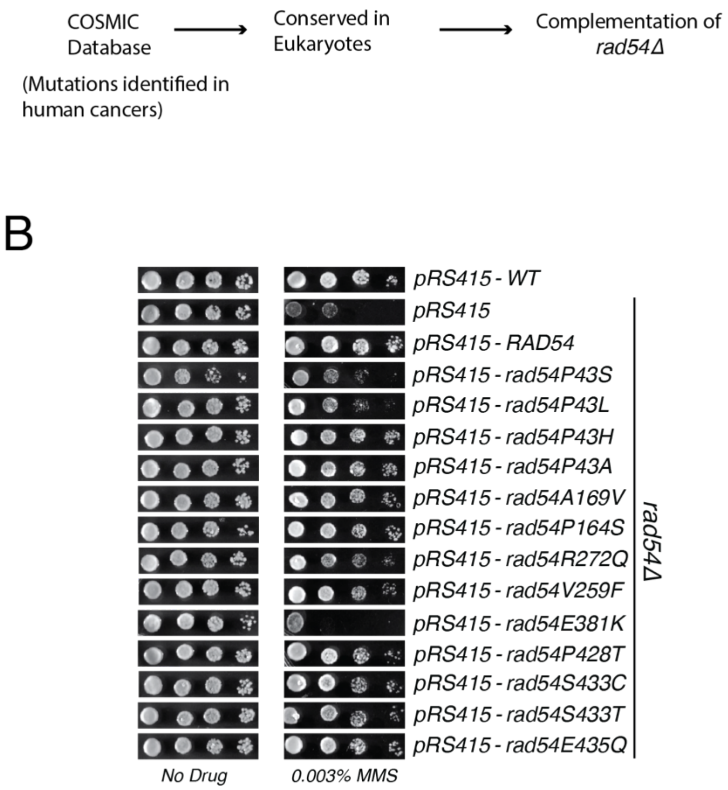


**Supplemental Figure 1: Identification of Rad54 reduction of function allele**
**(A).** Experimental set-up for identification of residues to test. Amino Acid substitutions were identified in human cancers and were tested to determine if they were conserved in eukaryotes. **(B).** Serial dilution yeast spot assay was used to test the complementation of *rad54*Δ sensitivity to the DNA alkylating agent MMS.

### Supplemental Figure 2

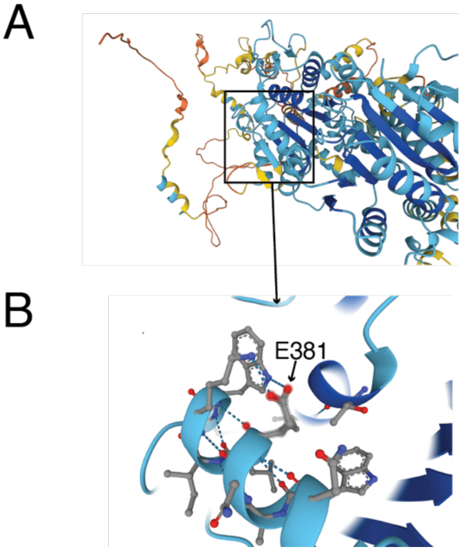


**Supplemental Figure 2: Rad54E381K likely destabilizes an essential helix**
**(A).** Alphafold2 structure of yeast Rad54. The alpha helix that contains the E381 residue is enclosed in a box. **(B).** Zoomed in view of the E381 residue. The residue likely contacts a Lysine and Histidine that stabilizes the helix. Conversion of the residue to Lysine disrupts the interaction and destroys the helix.

### Supplemental Figure 3

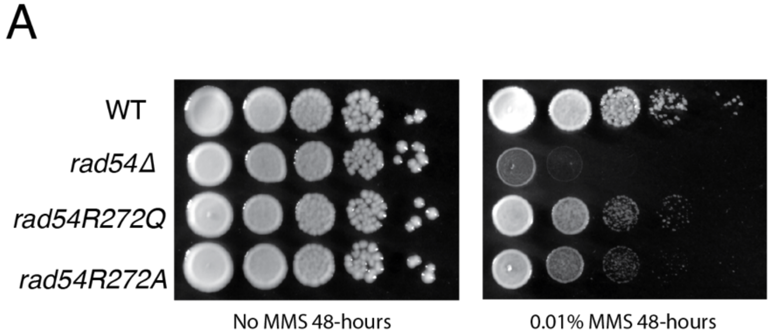


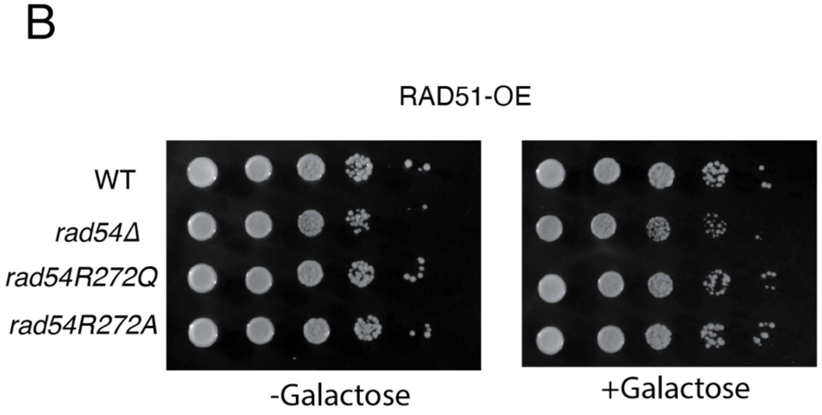


**Supplemental Figure 3: Rad54 mutants complement the Rad51 overexpression phenotype (A).** Serial dilution spot assay of WT, *rad54*Δ, *rad54R272Q, or rad54R272A* mutants spotted on YPD+/- 0.1% MMS. The spots represent a ten-fold dilution from the previous spot. **(B).** Serial dilution spot assay of WT, *rad54*Δ*, rad54R272Q,* or *rad54R272A* spotted on plates with or without galactose. The spots represent ten-fold dilutions from the previous spot.

### Supplemental Figure 4

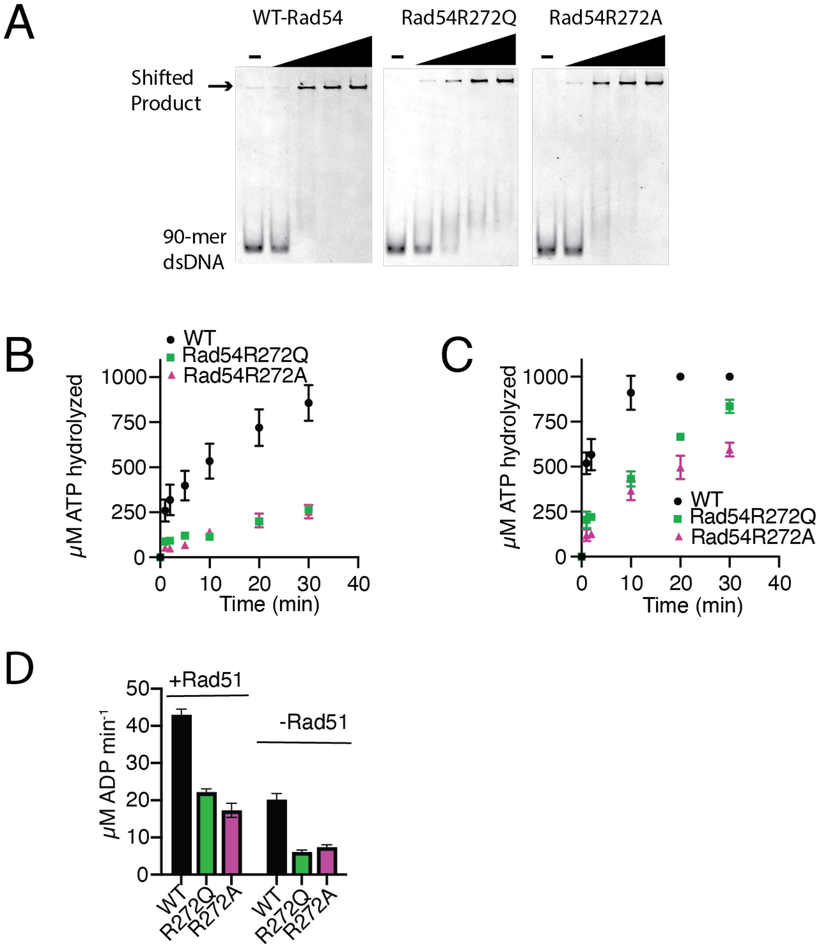


**Supplemental Figure 4: Rad54 mutants are active for ATP hydrolysis with and without Rad51**

**(A).** Electromobility shift assay (EMSA) for Rad54 binding to dsDNA. The Rad54, Rad54R272Q, and Rad54R272A concentrations are 0, 6.25, 12.5, 25 and 50 nM. **(B).** ATP hydrolyzed (µM) by WT (Black), Rad54R272Q (Green), and Rad54R272A (Magenta) without Rad51 as a function of reaction time. The error bars represent the standard deviation of three independent experiments. **(C).** ATP hydrolyzed (µM) by WT (Black), Rad54R272Q (Green), and Rad54R272A (Magenta) with Rad51 as a function of reaction time. The error bars represent the standard deviation of three independent experiments. **(D).** The rate of ATP hydrolysis (µM ADP formed $min^{-1}$) was measured from the linear portion of the graphs in A and B for WT (Black), Rad54R272Q (Green), and Rad54R272A (Magenta). The error bars represent the confidence in the fit of the data.

### Supplemental Figure 5

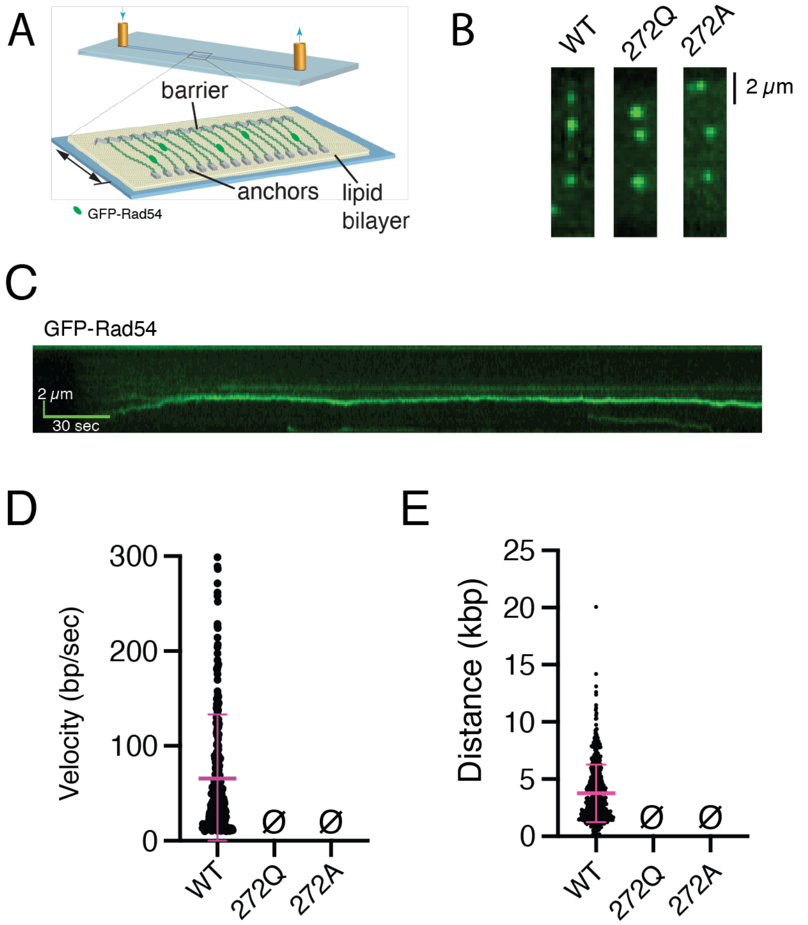


**Supplemental Figure 5: Rad54R272Q and Rad54R272A interact with dsDNA**
**(A).** Schematic cartoon illustrating the DNA curtains set-up to test Rad54 motor function dsDNA. **(B).** Widefield TIRFM microscope image of GFP-RAD54, GFP-Rad54R272Q, and GFP-Rad54R272A bound to dsDNA. **(C).** Representative kymograph illustrating movement of WT-Rad54 along dsDNA **(D).** The dot plot represents the velocities (N=278) for WT Rad54, Rad54R272Q, and Rad54R272A. The crossbar represents the mean of the data, and the error bars represent the standard deviation of the data. **(E).** The dot plot represents the distance traveled (N=584) for WT Rad54, Rad54R272Q, and Rad54R272A. The crossbar represents the mean of the data, and the error bars represent the standard deviation of the data.

### Supplemental Figure 6

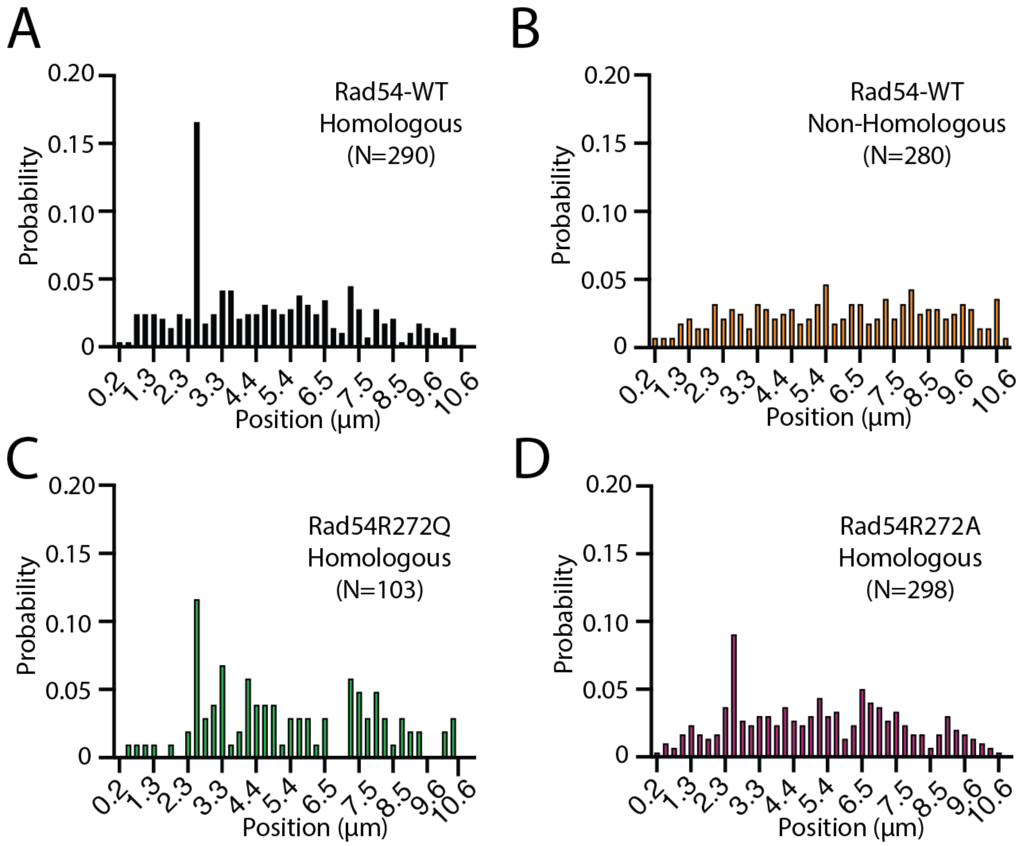


**Supplemental Figure 6: Rad54R272Q and Rad54R272A have a reduction in translocation and sequence alignment**

**(A).** Distribution of binding site probability for WT with homologous ssDNA (N=290). The data are also shown in Figure 3BC. **(B).** Distribution of critical site probability for WT with non-homologous ssDNA (N=290). The data are also presented in Figure 3BC. **(C).** Distribution of critical site probability for Rad54R272Q with homologous ssDNA (N=103). The data are also shown in Figure 3BC. **(D).** Distribution of binding site probability for Rad54R272A with homologous ssDNA (N=298). The data are also presented in Figure 3BC.

### Supplemental Figure 7

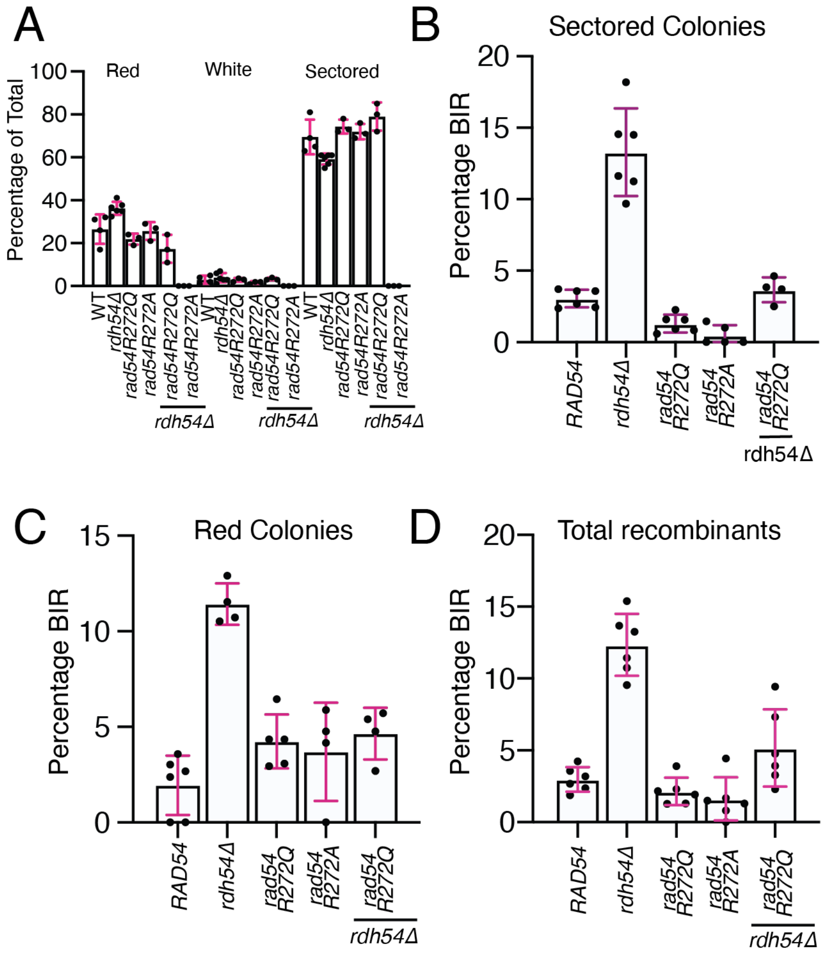


**Supplemental Figure 7: Rad54 mutants result in altered BIR outcomes**

**(A).** Measured outcomes for gene conversion tracts as observed for red, white, and sectored colonies. Strains assayed include WT, rdh54Δ, rad54R272Q, rad54R272A, rad54R272Q rdh54Δ, and rad54R272A rdh54Δ. **(B).** Graph comparing the percentage of BIR outcomes in sectored colonies only for WT, *rdh54Δ, rad54R272Q, rad54R272A, rad54R272Q rdh54Δ, and rad54R272A rdh54Δ*. The error bars represent the standard deviation for at least three independent experiments **(C).** Graph comparing the percentage of BIR-like outcomes in solid red colonies only for WT, *rdh54Δ, rad54R272Q, rad54R272A, rad54R272Q rdh54Δ, and rad54R272A rdh54Δ*. The error bars represent the standard deviation for at least three independent experiments **(D).** Graph representing the prevalence of BIR-like events in total recombinants for WT, *rdh54Δ, rad54R272Q, rad54R272A, rad54R272Q rdh54Δ, and rad54R272A rdh54Δ*.The error bars represent the standard deviation of at least three independent experiments

### Supplemental Information
### Supplemental Table 1

| Strains | Genotype | Source or reference |
|---|---|---|
| *LSY-2202-15D (WT)* | *MATa ade2-n his3::NatMX4 met22:klURA3* | *(Mazon et al. 2010)* |
| *LSY-2205-11C (WT)* | *MATalpha ade2-I lys2:GAL-ISCE1 his3:HphMX4* | *(Mazon et al. 2010)* |
| *BY4741 (WT)* | | *Dharmacon* |
| *BY4741* | *rad54Δ* | *Dharmacon* |
| *BY4741* | *RAD54::rad54R272Q* | *This study* |
| *BY4741* | *RAD54::rad54R272A* | *This study* |
| *JBC0105* | *15D-RAD54-KANMX* | *This study* |
| *JBC0106* | *11C-RAD54-KANMX* | *This study* |
| *JBC0107* | *15D-rad54R272Q-KANMX* | *This study* |
| *JBC0108* | *11C-rad54R272Q-KANMX* | *This study* |
| *JBC0109* | *15D-rad54R272A-KANMX* | *This study* |
| *JBC0110* | *11C-rad54R272A-KANMX* | *This study* |
| *JBC0241* | *15D-RAD54-KANMX rdh54::HIS3* | *This study* |
| *JBC0236* | *11C-RAD54-KANMX rdh54::HIS3* | *This study* |
| *JBC0243* | *15D-rad54R272Q-KANMX rdh54::HIS3* | *This study* |
| *JBC0238* | *11C-rad54R272Q-KANMX rdh54::HIS3* | *This study* |
| *JBC0245* | *15D-rad54R272A-KANMX rdh54::HIS3* | *This study* |
| *JBC0240* | *11C-rad54R272A-KANMX rdh54::HIS3* | *This study* |
| *JBC0210* | *S. cerevisiae protease deficient strain* | |
| ***Diploids*** | | *This study* |
| *JBC0105xJBC0106* | *RAD54-KANMX/RAD54-KANMX* | *This study* |
| *JBC0107xJBC0108* | *rad54R272Q/rad54R272Q* | *This study* |
| *JBC0109XJBC0110* | *rad54R272A/rad54R272A* | *This study* |
| *JBC0241xJBC0236* | *RAD54-KANMX/RAD54-KANMX rdh54/rdh54* | *This study* |
| *JBC243xJBC248* | *rad54R272Q/rad54R272Q rdh54/rdh54* | *This study* |
| *JBC0245XJBC0240* | *rad54R272A/rad54R272A rdh54/rdh54* | *This study* |

### Supplemental Table 2

| Name | Source |
|---|---|
| *pRS415* | |
| *pRS415-RAD54* | (Crickard et al., 2020) |
| *pRS415-rad54P43S* | This study |
| *pRS415-rad54P43L* | This study |
| *pRS415-rad54P43H* | This study |
| *pRS415-rad54P43A* | This study |
| *pRS415-rad54A169V* | This study |
| *pRS415-rad54P164S* | This study |
| *pRS415-rad54R272Q* | This study |
| *pRS415-rad54V259F* | This study |
| *pRS415-rad54E381K* | This study |
| *pRS415-rad54P428T* | This study |
| *pRS415-rad54S433C* | This study |
| *pRS415-rad54S433T* | This study |
| *pRS415-rad54E435Q* | This study |
| *pRS415-rad54R272A* | This study |
| *pRS415-rad54Y562A* | This study |
| *pRS415-rad54D769A* | This study |
| *pRS415-rad54D769H* | This study |
| pRS305-*RAD54-KANMX* | This study |
| pRS305-*rad54R272Q-KANMX* | This study |
| pRS305-*rad54R272A-KANMX* | This study |
| pYES-GST-RAD54 | |
| pYES-GST-GFP-RAD54 | Crickard et al. 2018 |
| pYES-GST-rad54R272Q | This study |
| pYES-GST-GFP-rad54R272Q | This study |
| pYES-GST-rad54R272A | This study |
| pYES-GST-GFP-rad54R272A | This study |
| pYES-RAD51 | |
| pET11C-SUMO-yRad51 | Crickard et al. 2020 |
| P11D-sctRPA | |
| P11D-sctRPA-mCherry-70 | |
| pUC19 plasmid | New England Biolabs |

### Supplemental Table 3

| hRAD54L mutation | yRad54 residue |
|---|---|
| P50S | 43 |
| P50L | 43 |
| P50H | 43 |
| P50A | 43 |
| P112S | 164 |
| A117V | 169 |
| V141F | 259 |
| R154Q | 272 |
| E229K | 381 |
| P269T | 428 |
| S274C | 433 |
| S274T | 433 |
| E276Q | 435 |
| D610H* | 769 |

*The D610H mutation was also found in COSMIC, but the respective yeast mutant *rad54D769H* was not created for the initial complementation spot assay. This mutant was created for the targeted complementation spot assay after it was found that residue D769 is involved in a stabilizing interaction.

#### Supplemental Table 4
#### Gene Conversion outcomes

| Strain | Solid Red | Solid White | Sectored | Total |
|---|---|---|---|---|
| WT | 249 | 26 | 707 | 979 |
| *rad54R272Q/rad54R272Q* | 226 | 15 | 667 | 908 |
| *rad54R272A/rad54R272A* | 131 | 15 | 595 | 741 |
| *rdh54/rdh54* | 183 | 17 | 443 | 644 |
| *rad54R272Q/rad54R272Q rdh54Δ* | 178 | 19 | 291 | 488 |
| *rad54R272A/rad54R272A rdh54Δ* | 0 | 0 | 0 | 0 |

#### Supplemental Table 5
#### Sectored

| Strain | CO | NCO | BIR | Total | Chromosome Loss |
|---|---|---|---|---|---|
| WT | 356 | 329 | 22 | 707 | 0 |
| *rad54R272Q/rad54R272Q* | 401 | 260 | 6 | 667 | 0 |
| *rad54R272A/rad54R272A* | 354 | 237 | 4 | 595 | 3 |
| *rdh54/rdh54* | 202 | 187 | 54 | 443 | 0 |
| *rad54R272Q/rad54R272Q rdh54* | 224 | 52 | 15 | 291 | 5 |
| *rad54R272A/rad54R272A rdh54* | 0 | 0 | 0 | 0 | N/A |

#### Supplemental Table 6
#### Solid Red

| Strain | CO | NCO | BIR | Total |
|---|---|---|---|---|
| WT | 34 | 208 | 7 | 249 |
| *rad54R272Q/rad54R272Q* | 44 | 175 | 7 | 226 |
| *rad54R272A/rad54R272A* | 46 | 123 | 5 | 131 |
| *rdh54/rdh54* | 20 | 146 | 17 | 183 |
| *rad54R272Q/rad54R272Q rdh54* | 46 | 124 | 8 | 178 |
| *rad54R272A/rad54R272A rdh54* | 0 | 0 | 0 | 0 |

### Supplemental Table 7

| Name | Sequence | Homologous region |
|---|---|---|
| Labeling Oligo | Atto647N-CCGCCTCGCAGAACGGGCATTCCCT | N/A |
| 65-mer pUC19 homology | AGGGAATGCCCGTTCTGCGAGGCGGTGGATCTCAACAGCGGTAAGA TCCTTGAGAGTTTTCGCCCCGAAGAACGTTTTCCAATGATGAGC | 2348-2373 |
| 90-mer pUC19 homology | AGGGAATGCCCGTTCTGCGAGGCGGGGTGCACGAGTGGGTTACATCG AACTGGATCTCAACAGCGGTAAGATCCTTGAGAGTTTTCGCCCCG AAGAACGTTTTCCAATGATGAGC | 2283-2373 |
| 130-mer pUC19 homology | AGGGAATGCCCGTTCTGCGAGGCGGGGTGCACGAGTGGGTTACATCG AACTGGATCTCAACAGCGGTAAGATCCTTGAGAGTTTTCGCCCCGAAG AACGTTTTCCAATGATGAGCACTTTTAAAGTTCTGCTATGTGGCGCGGT ATTATCCCGTA | 2243-2373 |
| 90-mer Lambda homology | Atto647N-GATGTTCTGCTGGATATGCACTTTTCCGGGC TGACGTACACCGTGCTCAGCCTGTTTTTCA GCGATCCGGATATGCATCCGCTGGATTTC | 10,253-10,342 |
| 90-mer Lambda | | N/A |
| EMSA Unlabeled strand | GACCATGATTACGCCAAGCTTGCATGCCTGCAGGTCGACTCTAGAGGATC CCCGGGTACCGAGCTCGAATTCACTGGCCGTCGTTTTACA | |
| 90-mer-fluor EMSA | Atto647N- TGTAAAACGACGGCCAGTGAATTCGAGCTCGGTACCCGGGGATCCTCTAG AGTCGACCTGCAGGCATGCAAGCTTGGCGTAATCATGGTC | |
